# Histone H3 threonine 11 phosphorylation is catalyzed directly by the meiosis-specific kinase Mek1 and provides a molecular readout of Mek1 activity *in vivo*

**DOI:** 10.1101/106245

**Authors:** Ryan Kniewel, Hajime Murakami, Yan Liu, Masaru Ito, Kunihiro Ohta, Nancy M. Hollingsworth, Scott Keeney

**Author notes:** Present address: Department of Environmental Biology, Centro de Investigaciones Biológicas, Consejo Superior de Investigaciones Científicas (CIB-CSIC), Madrid, Madrid, Spain. Present address: OCIO, Northwell Health System, New Hyde Park, New York, USA. Present address: Department of Microbiology and Molecular Genetics, University of California, Davis, USA. Equal contribution (Ryan Kniewel, Hajime Murakami).

## Abstract

*Saccharomyces cerevisiae* Mek1 is a CHK2/Rad53-family kinase that regulates meiotic recombination and progression upon its activation in response to DNA double-strand breaks (DSBs). The full catalog of direct Mek1 phosphorylation targets remains unknown. Here, we show that phosphorylation of histone H3 on threonine 11 (H3 T11ph) is induced by meiotic DSBs in *S. cerevisiae* and *Schizosaccharomyces pombe*. Molecular genetic experiments in *S. cerevisiae* confirmed that Mek1 is required for H3 T11ph and revealed that phosphorylation is rapidly reversed when Mek1 kinase is no longer active. Reconstituting histone phosphorylation *in vitro* with recombinant proteins demonstrated that Mek1 directly catalyzes H3 T11 phosphorylation. Mutating H3 T11 to nonphosphorylatable residues conferred no detectable defects in otherwise unperturbed meiosis, although the mutations modestly reduced spore viability in certain strains where Rad51 is used for strand exchange in place of Dmc1. H3 T11ph is therefore mostly dispensable for Mek1 function. However, H3 T11ph provides an excellent marker of ongoing Mek1 kinase activity *in vivo*. Anti-H3 T11ph chromatin immunoprecipitation followed by deep sequencing demonstrated that H3 T11ph was highly enriched at presumed sites of attachment of chromatin to chromosome axes, gave a more modest signal along chromatin loops, and was present at still lower levels immediately adjacent to DSB hotspots. These localization patterns closely tracked the distribution of Red1 and Hop1, axis proteins required for Mek1 activation. These findings provide insight into the spatial disposition of Mek1 kinase activity and the higher order organization of recombining meiotic chromosomes.

**bioRxiv version 2 (June 2017):** One major experimental change was incorporated into the revised manuscript: We repeated the anti-H3 T11ph ChIP-seq experiment on larger scale, including two meiotic time points from each of two wild type cultures and one time point from a spo11-Y135F mutant culture. To facilitate comparison of different samples, we used meiotic S. pombe cells as a spike-in control for all samples for both anti-H3 and anti-H3 T11ph ChIP-seq. Most conclusions described in the first bioRxiv submission were confirmed, but the improved datasets allowed us to derive more detailed information in particular about H3 T11ph patterns around DSB sites.

**bioRxiv version 3 (October 2017):** The following experimental changes were incorporated, along with more minor changes in response to reviewer comments:

- We added previously unpublished ChIP-seq data for Red1 protein, generated by Masaru Ito and Kunihiro Ohta, who have been added as coauthors.
- We repeated key experiments with the *H3-T11V* single point mutant. No conclusions were changed relative to prior experiments with the *H3-S10, T11V* mutant.
- We repeated the analysis of spore viability in a *dmc1 rad54-T132A* background using a more appropriate isogenic control, and recapitulated the original conclusion that the *H3-T11V* mutation modestly decreases spore viability in this sensitized background.

## INTRODUCTION

Meiotic recombination initiates with DNA double-strand breaks (DSBs) made by the topoisomerase-like transesterase Spo11 (Lam and Keeney 2014). DSBs occur throughout the genome, often but not always in hotspots that in *Saccharomyces cerevisiae* mostly overlap with nucleosome-depleted gene promoters (Ohta *et al.* 1994; Wu and Lichten 1994; Baudat and Nicolas 1997; Pan *et al.* 2011). Repair of meiotic DSBs by recombination helps form physical connections between homologous chromosomes that allow the chromosomes to segregate accurately at the first meiotic division (Hunter 2015). Because recombination defects can lead to mutations and/or aneuploidy, meiotic DSB repair is highly regulated (Subramanian and Hochwagen 2014; Hunter 2015).

A critical component of this regulation in yeast is Mre4/Mek1, a meiosis-specific paralog of the Rad53 checkpoint effector kinase (Rockmill and Roeder 1991; Leem and Ogawa 1992). In response to Spo11-generated DSBs, the kinases Tel1 and/or Mec1 (orthologs of mammalian ATM and ATR, respectively) become activated and phosphorylate the chromosome axis-associated protein Hop1 among other substrates (Carballo *et al.* 2008; Cheng *et al.* 2013; Penedos *et al.* 2015). The FHA (Forkhead-associated) domain of Mek1 then binds phosphorylated Hop1, resulting in Mek1 recruitment to chromosome axes where Mek1 undergoes activation (involving trans-autophosphorylation on T327 in its activation loop) and stabilizes Hop1 phosphorylation via positive feedback (Niu *et al.* 2005; Niu *et al.* 2007; Carballo *et al.* 2008; Chuang *et al.* 2012; Penedos *et al.* 2015). Activated Mek1 promotes inter-homolog bias in recombination, that is, the preferential use of a homologous chromosome rather than sister chromatid as the template for DSB repair (Niu *et al.* 2005; Carballo *et al.* 2008; Goldfarb and Lichten 2010; Kim *et al.* 2010; Hong *et al.* 2013; Lao *et al.* 2013; Subramanian *et al.* 2016). Mek1 does so in part by phosphorylating the Rad54 protein on threonine 132 (T132) (Niu *et al.* 2007; Niu *et al.* 2009). Rad54 is a member of the Swi2/Snf2 DNA-dependent-ATPase chromatin remodeling family and is a binding partner of the strand exchange protein Rad51 (Heyer *et al.* 2006). Mek1-dependent phosphorylation of Rad54 attenuates the interaction with Rad51, allowing the meiosis-specific strand exchange protein Dmc1 to predominate (Niu *et al.* 2009). Mek1 also directly phosphorylates the T40 residue of Hed1; this stabilizes the Hed1 protein and thereby promotes its function as a negative regulator of Rad51 strand exchange activity (Callender *et al.* 2016). Additionally, Mek1 promotes the repair of interhomolog strand invasion intermediates through a pathway required for chromosome synapsis and the generation of crossovers whose distribution shows interference (Chen *et al.* 2015). Finally, *MEK1* is required for checkpoint arrest or delay of meiotic progression in response to unrepaired DSBs (Lydall *et al.* 1996; Xu *et al.* 1997).

The full array of direct Mek1 phosphorylation substrates remains unknown, as only three direct targets have been definitively proven thus far: Mek1 itself, Rad54, and Hed1 (Niu *et al.* 2007; Niu *et al.* 2009; Callender *et al.* 2016). Additional Mek1-dependent phospho-proteins have been identified by mass spectrometry and other approaches, including T11 of histone H3 (Govin *et al.* 2010; Suhandynata *et al.* 2016). However, a number of Mek1-dependent phosphorylation events are known or suspected to be indirect (Suhandynata *et al.* 2016). For example, Mek1 is required for phosphorylation of the synaptonemal complex protein Zip1, but the kinase directly responsible is Cdc7-Dbf4, not Mek1 (Chen *et al.* 2015). Moreover, H3 T11 phosphorylation has been reported as being catalyzed in vegetative cells by other kinases [the pyruvate kinases Pyk1 and, to a lesser extent, Pyk2 (Li *et al.* 2015)], which could in principle be regulated by Mek1 in meiosis. Therefore, whether H3 T11 is a direct substrate for Mek1 remains to be established.

Mek1 activity plays out in the context of elaborate higher order chromosome structures. Early in meiotic prophase, sister chromatids form co-oriented arrays of DNA loops that are anchored along a linear proteinaceous axis (Zickler and Kleckner 1999; Kleckner 2006). Prominent components of these axes include sister chromatid cohesion proteins (including the meiosis-specific Rec8 subunit), Mek1, Hop1, and another meiosis-specific chromosome structural protein, Red1 (Smith and Roeder 1997; Bailis and Roeder 1998; Klein *et al.* 1999; Panizza *et al.* 2011).

In cytological experiments, immunostaining foci of recombination proteins are axis-associated, indicating that recombination occurs in proximity to axes (reviewed in Zickler and Kleckner 2015). However, there is an anticorrelation between the DNA sequences preferentially bound by axis proteins (Rec8, Hop1, Red1) and the DNA sequences that often experience Spo11-induced DSBs, which suggests that recombination usually involves the DNA in chromatin loops rather than the DNA embedded in axes (Gerton *et al.* 2000; Blat *et al.* 2002; Pan *et al.* 2011; Panizza *et al.* 2011). To reconcile this paradox, the “tethered-loop/axis complex” (TLAC) model proposes that DNA segments residing on chromatin loops incur DSBs but are recruited, or tethered, to axes by interactions between recombination proteins and axis proteins (Kleckner 2006; Panizza *et al.* 2011). The TLAC model provides a framework for understanding spatial organization of recombining chromosomes, but there is as yet little direct molecular data demonstrating the proposed functional interactions between axes and DSB sites.

How Mek1 fits into this proposed organization also remains unknown. Immunocytology suggests that Mek1 protein is localized primarily on axes (Bailis and Roeder 1998; Subramanian *et al.* 2016), supported by the dependence of Mek1 activity on axis proteins (Niu *et al.* 2007; Carballo *et al.* 2008). However, Mek1 exerts its known recombination-controlling activity (directly or indirectly) at sites of DSBs. The TLAC model can account for Mek1 acting at both places, but where Mek1 kinase activity actually occurs remains unexplored because of a lack of a molecular marker for the active kinase.

In this study we demonstrate that Mek1 directly phosphorylates histone H3 T11 in response to meiotic DSBs in *S. cerevisiae*. H3 T11ph is dispensable for Mek1 function during unperturbed meiosis, so the purpose of this phosphorylation remains unclear. Nevertheless, we demonstrate the utility of H3 T11ph as a direct molecular marker for active Mek1 by examining the genome-wide localization of H3 T11ph. Our findings suggest that Mek1 exerts its activity at axis association sites but also across chromatin loops.

## MATERIALS AND METHODS

### Strains and histone mutagenesis strategy

*S. cerevisiae* and *S. pombe* strains are listed in **Supplemental Table S1**. *S. pombe* strains were generously provided by G. Smith, Fred Hutchinson Cancer Research Center. Histone gene deletion strains and plasmids expressing H3 T11 mutants from Govin et al. (2010) were generously provided by S. Berger, University of Pennsylvania. *S. cerevisiae* strains were of the SK1 strain background. Because of concerns about effects of plasmid (in)stability on the ability to score phenotypes of histone mutants and to reliably measure meiotic parameters because of cell-to-cell heterogeneity within a culture (see Results), we opted to avoid plasmid shuffle systems that have been used by others (Ahn *et al.* 2005; Govin *et al.* 2010). Instead, strategies involving stable integration or gene replacement were employed, as follows.

#### Histone gene replacements

*S. cerevisiae* histone genes are arranged in divergently oriented pairs expressing either H3 and H4 or H2A and H2B; there are two of each pair, i.e., two copies encoding each histone. The S10A and T11V mutations were introduced into plasmid-borne copies of *HHT1* and *HHT2* by QuikChange site-directed mutagenesis (Agilent Technologies). These mutant alleles were then introduced sequentially into SK1 strain SKY165 by one-step gene replacements using DNA fragments containing ≥270 bp arms of homology. Targeting constructs included selectable drug resistance markers: *kanMX4* ∼366 bp downstream of the *HHT1* ORF and *hphMX4* ∼250 bp downstream of *HHT2*.

#### Stable integration of histone gene cassettes

A histone cassette integration strategy was employed using pRS305-based plasmids (Sikorski and Hieter 1989) integrated into the *leu2::hisG* locus. Integrations were performed to try to maintain balanced gene dosage for the four core histones. The parental strain for the H2A/H2B/H3/H4 histone cassette integrations was created in a multistep process by first transforming a pRS316-based *URA3* histone cassette covering plasmid containing a single copy of each histone gene (pRK12; *HTA1-HTB1*, *HHT2-HHF2*) into diploid SKY165. Next, the histone gene pairs, *HHT2-HHF2* and *HTA1-HTB1* (which are required for proper meiosis (Norris and Osley 1987)), were deleted sequentially and replaced with the *hphMX* and *natMX* markers, respectively. The deletions were confirmed by Southern blot and the strain was sporulated to yield a Ura^+^, Nat^R^, Hyg^R^, *MATα* haploid. A second *MAT***a** haploid strain was created by sequentially deleting the other (non-essential) histone gene pairs, *HTA2-HTB2* and *HHT1-HHF1*, which were replaced by the *kanMX* and *natMX* markers, respectively, and confirmed by Southern blot. These two haploids were mated to form a compound heterozygote, then tetrads were dissected and resulting haploids carrying all four histone gene-pair deletions were mated to form a histone integration host strain (SKY2283) with the genotype: *hht1-hhf1*Δ::*kanMX/”*, *hht2-hhf2*Δ::*natMX/”, hta1-htb1*Δ::*hphMX/”, hta2-htb2*Δ::*natMX/”, pRK12[CEN/ARS, URA3, HTA1-HTB1, HHF2-HHT2]*.

A parental strain for the H3/H4 histone cassette integrations was created by dissecting tetrads from the *hht2-hhf2*Δ::*natMX/”, pRK12* strain described above prior to deletion of *HTA1-HTB1*. This dissection yielded a Ura^+^, Nat^R^, *MAT***a** haploid that was crossed with the second haploid strain described above (*hta2-htb2*Δ::*natMX, hht1-hhf1*Δ::*kanMX*). Tetrad dissection yielded *MAT***a** and *MATα* haploid progeny (SKY3166 and SKY3167, respectively) with the following genotype: *hht1-hhf1*Δ::*kanMX, hhf2-hht2*Δ::*natMX, hta2-htb2*Δ::*natMX, pRK12.*

All histone mutant integration constructs were created by QuikChange site-directed mutagenesis. The first was a H3/H4 replacement using a pRS305-based plasmid (pRK77) containing *LEU2*, *HHT2*-*HHF2* that was linearized by AflII digestion to target integration to *leu2::hisG* and transformed into haploids SKY3166 and SKY3167. The second was an H2A/H2B/H3/H4 replacement using a pRS305-based plasmid (pRK24) containing *LEU2*, *HTA1-HTB1* and *HHF2-HHT2* that was linearized by AflII digestion and transformed into diploid SKY2283. In both cases, the core-histone covering plasmid pRK12 was counterselected by growth on 5-fluoroorotic acid (FOA). Colony PCR of Leu+, Ura– transformants was used to verify the proper integration into the *leu2::hisG* locus using primer sets flanking both junctions as well as verification of the mutations in *hta1* and *hht2* by engineered restriction enzyme site polymorphisms and/or sequencing. In the case of the SKY3166/3167 transformants, haploid integrants were subsequently mated to create diploids. SKY2283 hemizygous integrants were sporulated to produce haploid progeny that were then mated to create homozygous diploids.

### *S. cerevisiae* and *S. pombe* cultures

*S. cerevisiae* was cultured at 30°C with asynchronous vegetative (cycling) cultures in YPD (1% yeast extract, 2% peptone, 2% dextrose). Camptothecin treatment (20 μM) was performed for 2 hr at 30° in 250 ml flasks shaking at 250 rpm in 10 ml cultures of SKY165 at an initial cell density of ∼9 × 10^7^ cells/ml. An untreated culture was incubated in parallel, while a separate 10 ml aliquot in a vented T-75 flask was exposed to X-rays for 60 min at room temperature using an X-RAD 225C X-ray irradiator (Precision X-ray, Inc.) corresponding to a dose of 400 Gy. Alternatively, 10 ml of culture at ∼7 × 10^7^ cells/ml was exposed to X-rays for 60 min on ice, with untreated cells also held on ice. With both exposure conditions, cells were subsequently allowed to recover at 30°, shaking at 225 rpm for 60 min (room temperature exposure) or 30 min (exposure on ice) before fixing in 20% trichloroacetic acid (TCA), pelleting and storage at -80° until extract preparation.

For inhibition of Mek1-as *in vivo*, an SKY3095 culture was divided equally four hours after transfer to sporulation medium and 10 μl 100% DMSO was added to half while the other received 1 μM final concentration of 1-NA-PP1 (1-(1,1-Dimethylethyl)-3-(1-naphthalenyl)-1Hpyrazolo[3,4-d]pyrimidin-4-amine) dissolved in DMSO (Wan *et al.* 2004). The return-to-growth recombination assays using *arg4* heteroalleles were carried out in triplicate as described (Martini *et al.* 2006). Pulsed-field gel electrophoresis (PFGE) and Southern blotting on DNA from meiotic cultures prepared using the SPS method was performed as described (Murakami *et al.* 2009). Plasmid shuffling and meiotic cultures using plasmids and the SK1 histone gene deletion strain obtained from S. Berger were carried out as described (Govin *et al.* 2010).

*S. pombe* haploid *pat1-114* sporulation was carried out as described (Hyppa and Smith 2009). For *S. cerevisiae* meiotic cultures, strains were thawed on YPG plates (1% yeast extract, 2% peptone, 3% glycerol, 2% agar) and incubated for ∼2 days, then streaked for single colonies on YPD plates and grown ∼2 days. Single diploid colonies were inoculated in 5 ml YPD and grown overnight. Cultures were diluted in YP+1% potassium acetate presporulation medium to ∼1.2 × 10^6^ cells/ml, grown for 13.5 hours at 225 rpm for ChIP and 250 rpm for all other experiments. Cells were pelleted, washed in sterile water and resuspended in the same preculture volume of 2% potassium acetate to a density of ∼2–3 × 10^7^ cells/ml. This corresponds to 0 hr of the meiotic time course. Sporulation was at 225 rpm for chromatin immunoprecipitation (ChIP) and 250 rpm for all other experiments. Unless indicated otherwise, statistical significance of spore viabilities was assessed by Fisher’s exact test, treating dissected tetrads as a random spore population. Meiotic progression was assessed in culture aliquots fixed with 50% ethanol and stained with 5 μg/ml 4’,6-diamidino-2-phenylindole (DAPI).

### Whole-cell extracts and western blotting

Culture aliquots of OD_600_ = 10 for *S. pombe* or ∼3.2 × 10^8^ cells for *S. cerevisiae* were washed in 20% TCA, pelleted and stored at -80°C until ready for use. Aliquots were thawed, resuspended in 20% TCA and disrupted by bead beading at 4° using 0.5 mm zirconia/silica or glass beads and monitored microscopically until near complete disruption was observed. Samples were collected by centrifugation, then washed with 5% TCA and the pellet was resuspended in 1× NuPAGE LDS Sample Loading Buffer (Life Technologies Corp.) with 100 mM dithiothreitol (DTT). Samples were separated on 12% bis-Tris NuPAGE gels in 1× MOPS or MES running buffer (Life Technologies Corp.) or 15% Laemmli gels (Laemmli 1970). Proteins were blotted to polyvinyldifluoride (PVDF) membranes by semi-dry electrophoretic transfer using the iBlot system (Life Technologies Corp.) or in Tris-glycine (25 mM Tris base, 192 mM glycine, 10% methanol, 0.04% sodium dodecyl sulfate) at 100 mA constant for 70 min (TransBlot SD Transfer Cell, Bio-Rad Laboratories, Inc.). Membranes were air dried, then incubated with one of the following rabbit primary antibodies diluted in 5% non-fat milk (NFM) in Tris-buffered saline-Tween buffer (TBST; 25 mM Tris-HCl pH 7.4, 137 mM NaCl, 2.7 mM KCl, 0.1% Tween 20): anti-H3 polyclonal (Abcam 1791) diluted 1:10,000; anti-H3 T11ph mononclonal (EMD Millipore 05-789) diluted 1:1000; anti-H3 T11ph polyclonal (Active Motif 39151) diluted 1:1000; anti-H3 S10ph monoclonal (EMD Millipore 05-817) diluted 1:1000; anti-H3 S10ph polyclonal (EMD Millipore 06-560) diluted 1:1000; or anti-H2A S129ph polyclonal (Abcam 15083) diluted 1:500. The polyclonal secondary antibody used was horseradish peroxidase-conjugated goat anti-rabbit (Pierce/ThermoFisher Scientific 31462 or 31460) diluted 1:10,000 in TBST with visualization by the ECL-Plus kit (GE Healthcare Ltd.) exposed to chemiluminescent film or charged-coupled device (CCD) camera (Imagestation, Eastman Kodak Company).

### Validation of anti-phospho-H3 antibodies

Two commercial anti-H3 T11ph antibodies yielded Spo11-dependent bands at the expected size for H3 on western blots, but the monoclonal gave more robust signal with less background (**Figure 1B**). To more definitively characterize the specificity of these antibodies, we incubated them with synthetic peptide arrays containing different H3 modification states (Active Motif MODified histone peptide array)(**Supplemental Table S2**). The monoclonal anti-H3 T11ph antibody reacted strongly with all peptides containing T11ph regardless of other modifications present, unless S10 was also phosphorylated, in which case reactivity was strongly or completely lost (**Supplemental Figure S1Ai**). This monoclonal antibody was highly specific, as little to no cross-reactivity was observed for unmodified H3 peptides, H3 peptides carrying other modifications, or peptides from other histones, including peptides phosphorylated at other sites (H3 S10ph, H3 S28ph, H4 S1ph, H2A S1ph, H2B S14ph) (**Supplemental Figure S1Ai**). In a more limited analysis, the polyclonal anti-H3 T11ph antibody bound specifically to a peptide with trimethylated H3 K9 (K9me3) as well as T11ph, but not to unmodified or S10ph peptides from H3 or full-length unmodified histones (**Supplemental Figure S1B**). However, this polyclonal antibody showed substantial non-histone cross-reactivity against yeast whole-cell extracts that was not observed for the monoclonal anti-H3 T11ph antibody (**Figure 1B**).

**Figure 1.**
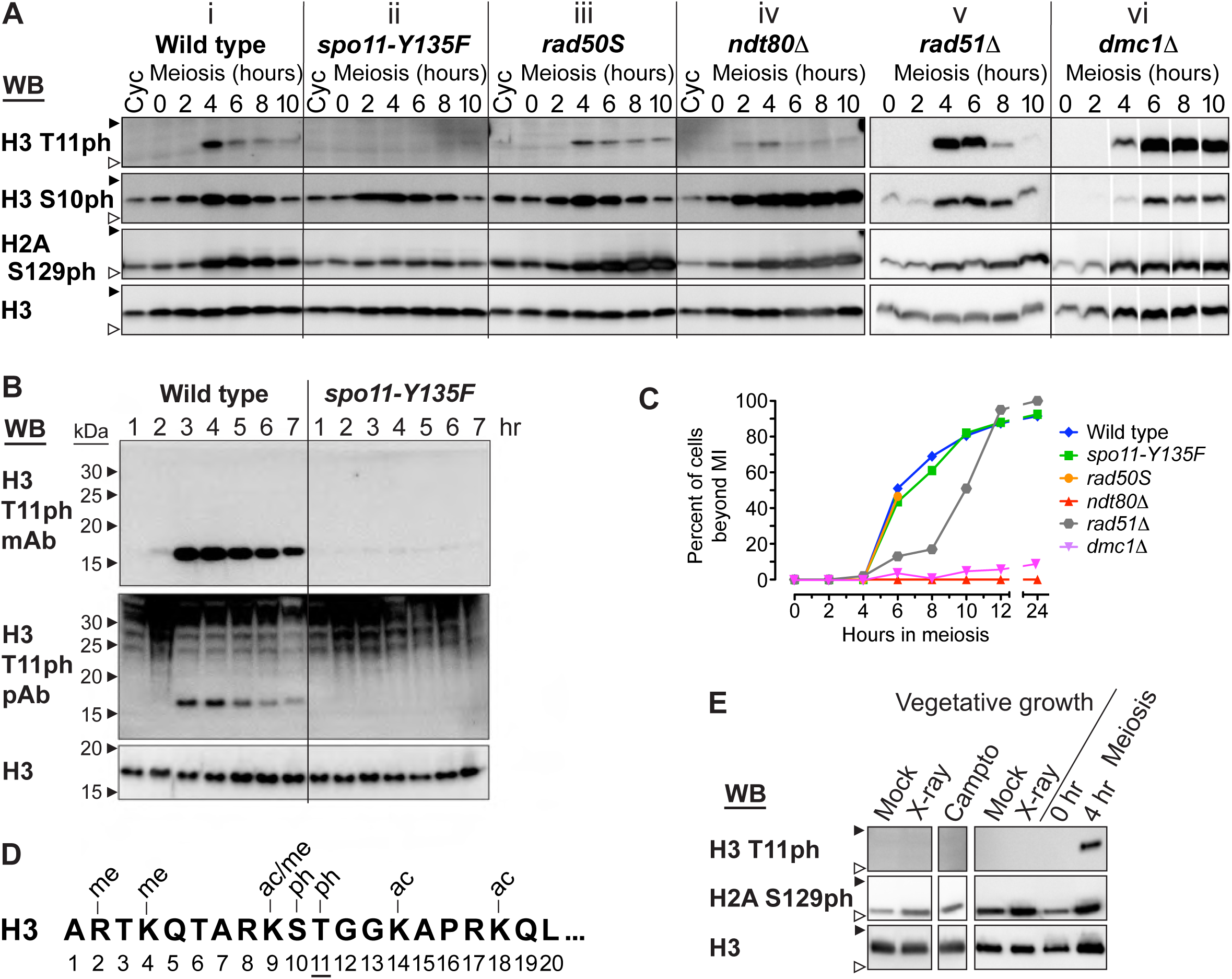
H3 T11 phosphorylation in *S. cerevisiae* meiosis. **(A)** Western blots of whole-cell extracts from asynchronous cycling vegetative (Cyc) and synchronized meiotic culture time points in wild-type and mutant strains. In panels i–iv, the antibodies used were anti-H3 T11ph polyclonal (Active Motif 39151), anti-H3 S10ph monoclonal (EMD Millipore 05-817), anti-H2A S129ph/γ-H2A (Abcam 15083), and anti-H3 (Abcam 1791). For panels v and vi, anti-H3 T11ph monoclonal (EMD Millipore 05-789) and anti-H3 S10ph polyclonal (EMD Millipore 06-560) were used; other antibodies were the same. Interstitial lanes were removed from the blot images in panel vi to match time points in other panels. Filled and open arrowheads indicate 20 and 15 kDa molecular weight markers, respectively. **(B)** Western blot comparison of anti-H3 T11ph monoclonal (mAb; EMD Millipore 05-789) and polyclonal (pAb; Active Motif 39151) antibodies. **(C)** Meiotic progression assessed by DAPI staining. Cells with ≥2 DAPI-staining bodies were scored as having progressed past the first meiotic division; n ≥ 100 cells per time point. The *rad50S* culture was not quantified past 6 hr because of nuclear fragmentation. **(D)** The first twenty amino acids in histone H3 and modifications known to occur in *S. cerevisiae* or *S. pombe*: ac, acetylation; me, methylation; ph, phosphorylation. **(E)** Meiosis-specificity of DNA damage-induced H3 T11ph. Asynchronous vegetative cultures of wild type were treated with genotoxins that induce DSBs, then whole-cell extracts were prepared and analyzed by western blotting for H3 T11ph. Cultures in the left panel were untreated (Mock) or treated with X-rays (400 Gy) or camptothecin (20 μM) at room temperature. An interstitial lane was deleted from the blot image for this panel. Cultures in the right panel were untreated or treated with X-rays (400 Gy) on ice. Premeiotic (0 hr) and meiotic (4 hr) cultures were included as controls. The anti-H3 T11ph monoclonal (EMD Millipore 05-789) was used. Arrowheads are as defined in panel A.

Both the monoclonal and the polyclonal anti-H3 S10ph antibodies we used reacted with phospho-S10 H3 peptide on dot blots, but with some background signal for full-length histone H3 (**Supplemental Figure S1B**). Similarly, the polyclonal anti-H3 S10ph antibody detected S10ph on the peptide array, including in the context of other nearby modifications, unless T11 was also phosphorylated (**Supplemental Figure S1Aii**). Again, however, modest cross-reactivity was seen with other histone H3 and H4 peptides, thus the anti-S10ph antibodies are less specific than the monoclonal anti-T11ph antibody.

### *In vitro* kinase assays

GST-Mek1 and GST-mek1-as were affinity purified on glutathione sepharose as described (Niu *et al.* 2009; Lo and Hollingsworth 2011).

#### Radiolabeling method

Reactions included 2 μg of recombinant *S. cerevisiae* histone H3 or 5 μg H3 1-20 peptides, 250 ng GST-Mek1, 0.4 mM ATP and 10 μCi [γ-^32^P]-ATP (6000 Ci/mmol; PerkinElmer, Inc.) in 25 μl total volume in a buffer containing 50 mM HEPES-NaOH pH 7.5, 150 mM NaCl, 10 mM MgCl_2_, 0.5 mM DTT and 1× each of Roche phosphatase and protease inhibitor cocktails. Reactions were incubated at 30° for 30 min then resolved on 12% bis-Tris NuPAGE gels in 1× MES running buffer and transferred to PVDF via the iBlot system or Coomassie stained and dried for autoradiography on a Fujifilm FLA 7000. Primary antibody was rabbit anti-H3 T11ph polyclonal (Active Motif 39151) diluted 1:500, with secondary antibody and detection carried out as described above.

#### Semi-synthetic epitope method

GST-Mek1-as target labeling and detection followed previously described methods (Niu *et al.* 2009; Lo and Hollingsworth 2011). Reactions included 2 μg of recombinant *S. cerevisiae* histone H3, 2 μg GST-Mek1 or 0.76 μg GST-Mek1-as, 0.4 mM ATPγS or 6-Fu-ATPγS (N^6^-furfuryladenosine-5’-O-3-thiotriphosphate, Axxora, LLC), and 0.2 mM ATP in 25 μl total volume in a buffer containing 50 mM Tris-HCl pH 7.5, 150 mM NaCl, 10 mM MgCl_2_ and 0.5 mM DTT. Reactions were incubated at 30° for 30 min, then *p-*nitrobenzyl mesylate (PNBM in DMSO, Abcam/Epitomics 3700-1) was added to 2.5 mM and incubated at room temperature for 90 min. Samples were electrophoresed on 4–12% bis-Tris NuPAGE gels in 1× MES running buffer, followed by semi-dry transfer to PVDF at 25 V constant for 60 min. Membranes were blocked in 5% NFM-TBST, primary antibodies were rabbit anti-thiophosphate ester monoclonal (Abcam/Epitomics 2686-1) diluted 1:5000 or rabbit anti-H3 T11ph monoclonal (EMD Millipore 05-789) diluted 1:1000, with secondary antibody and detection carried out as described above.

### Histone ChIP-sequencing

The ChIP-seq protocol was based on a previously described method (Wal and Pugh 2012). Two independent wild type (SKY165) and one *spo11-Y135F* (SKY198) meiotic cultures were prepared as described (Murakami and Keeney 2014) and 4 × 10^9^ cells were harvested at 3 and 4 hr (wild type), and 3.5 hr (*spo11-Y135F*) after the meiosis induction. Cells were fixed with 1% formaldehyde for 15 min at room temperature, with mixing at 50 rpm. Crosslinking was quenched by adding glycine to 131 mM for 5 min, cells were washed with water, resuspended in ice-cold ST buffer (10 mM Tris-HCl pH 7.4, 100 mM NaCl and 1× each of Roche phosphatase and protease inhibitor cocktails). To compare different samples, we used *S. pombe* cells as a spike-in control. *S. pombe* cells (SKY2594) harvested at 4.5 hr in meiosis were fixed and washed with the same condition described above. An aliquot of 4 × 10^7^ *S. pombe* cells (1% of the number of *S. cerevisiae* cells) were added to each sample.

Cells were resuspended in FA lysis buffer (50 mM HEPES-NaOH pH 7.5, 150 mM NaCl, 2 mM EDTA, 1% Triton X-100, 0.1% sodium deoxycholate, 10 μg/ml each of leupeptin, pepstatin A, and chymostatin, 1 mM PMSF, 1× each of Roche phosphatase and protease inhibitor cocktails) and disrupted using zirconia/silica beads (0.5 mm, Biospec Products, Inc. 11079105z) and a FastPrep-24 (MP Biomedicals) with 8 rounds of shaking at 6.5 m/s for 60 seconds. Lysates were pelleted by centrifugation at 15,000 rpm for 5 min at 4°, washed with NPS buffer (0.5 mM spermidine, 0.075% IGEPAL CA-630, 50 mM NaCl, 10 mM Tris-HCl pH 7.4, 10 mM MgCl^2^, 2 mM CaCl^2^, 10 μg/ml each of leupeptin, pepstatin A, and chymostatin, 1 mM PMSF, 1× each of Roche phosphatase and protease inhibitor cocktails) and resuspended in 3.6 mL NP-S buffer with 1 mM 2-mercaptoethanol. The resuspended pellet (chromatin) was solubilized by digestion with 25 units/ml of micrococcal nuclease (Worthington Biochemical Corp.) at 37° for 20 min. Digestion was terminated by adding EDTA to 10 μM and SDS to 0.05%. Chromatin was further solubilized by sonication (Biorupter Standard, Diagenode) on highest setting for two rounds of 30 sec with a 30 sec intervening rest. Solubilized chromatin was isolated by centrifugation at 16,000 rpm for 10 min at 4°, pooled and divided into two equal volumes.

Twenty μg of each antibody was added to the MNase-treated chromatin samples and incubated at 4° overnight on a rotisserie mixer [Antibodies: rabbit anti-H3 pAb (Abcam 1791); rabbit anti-H3 T11ph mAb (EMD Millipore 04-789)]. Immunoprecipitation was carried out by adding 200 μl protein G Dynabeads (Life Technologies Corp.) and incubating at 4° for 90 min on a rotisserie mixer. Beads were washed with 1 ml of the following buffers: NP-S buffer, FA lysis buffer, 2× FA high salt buffer (FA lysis buffer containing 1 M NaCl), 2× FA wash 2 buffer (FA lysis buffer containing 0.5 M NaCl), 2× FA wash 3 buffer (10 mM Tris-HCl pH 8, 250 mM LiCl, 2 mM EDTA, 1% IGEPAL, 1% sodium deoxycholate) and TE wash buffer (10 mM TrisHCl pH 8, 1 mM EDTA, 0.5% Triton X-100). Bound nucleosomes were eluted, reverse crosslinked, treated with RNase A and Proteinase K as described (Murakami and Keeney 2014). DNA was purified using PCR purification kit (Qiagen) and separated on a 1.5% agarose gel. Mononucleosome-sized DNA (∼150 bp) was extracted from the gel and prepared for 50 nt paired-end sequencing on the Hiseq platform (Illumina, Inc.) following standard Illumina protocols. Sequencing was performed at the Integrated Genomics Operation of Memorial Sloan-Kettering Cancer Center.

Paired-end 50 nt reads were mapped to the *S. cerevisiae* reference genome (sacCer2) and the Sanger Center’s *S. pombe* genome version of 7 August 2010 using BWA (version 0.7.12-r1039) MEM (Li 2013). Paired reads with an insert size more than 250 bp were filtered out and the rest were converted into coverage maps. All downstream analyses were carried out using R (<u>http://www.r-project.org/</u>) (R Development Core Team 2012). Each coverage map was normalized per 1000 reads from *S. pombe* chromosomes I and II. Cumulative S1-seq values were generated by calculating the cumulative sum of the top strand (or bottom strand) reads of S1-seq data (Mimitou *et al.* 2017) from the midpoint of each hotspot to 2 kb downstream (or upstream).

### Red1 ChIP-sequencing

The Red1 ChIP-seq protocol was based on a previously described method (Kugou *et al.* 2009). Meiotic cells (5 × 10^8^) expressing Flag-tagged Red1 (strain YKT190) were harvested at 3 hr after the meiosis induction. Cells were fixed with 1% formaldehyde for 10 min at room temperature and crosslinking was quenched by adding glycine to 125 mM for 5 min. Cells were washed with ice-cold Tris-buffered saline (TBS; 20 mM Tris-HCl pH 7.6, 150 mM NaCl).

Cells were resuspended in lysis 140 buffer (50 mM HEPES-KOH pH 7.5, 140 mM NaCl, 1 mM EDTA pH 8.0, 1% Triton X-100, 0.1% sodium deoxycholate, 1× Roche protease inhibitor cocktail) and disrupted using zirconia beads (0.5 mm, Yasui Kikai) and a Multi-Beads Shocker (Yasui Kikai) with four rounds of shaking at 2,300 rpm for 60 sec at 4°. Lysates were sonicated using Covaris (Covaris Inc.) to obtain genomic DNA fragments averaging 300 bp and the supernatant (whole cell extract (WCE)) was collected as after centrifugation at 15,000 rpm for 15 min at 4°.

Immunoprecipitation was carried out by adding 3 μl of anti-FLAG antibody (Wako) coupled to 80 μl of protein A Dynabeads (Life Technologies Corp.) and incubating at 4° for 3.5 hr by rotating. Beads were washed with 1 ml of the following buffers: lysis 140 buffer twice, lysis 500 buffer (50 mM HEPES-KOH pH 7.5, 500 mM NaCl, 1 mM EDTA pH 8.0, 1% Triton X-100, 0.1% sodium deoxycholate) once, LiCl/detergent buffer (10 mM Tris-HCl pH 8.0, 250 mM LiCl, 1 mM EDTA pH 8.0, 0.5% Nonident P-40, 0.5% sodium deoxycholate) twice, and TE (10 mM Tris-HCl pH 8.0, 1 mM EDTA pH 8.0) once. Bead-bound chromatin was eluted in 100 μl of TE/1% SDS (50 mM Tris-HCl pH 8.0, 10 mM EDTA pH 8.0, 1% SDS) by incubating at 65° for 15 min and then in 150 μl of TE/0.67% SDS (50 mM Tris-HCl pH 8.0, 10 mM EDTA pH 8.0, 0.67% SDS) by incubating at 65° for 5 min. For input, a small volume of WCE was mixed with TE/1.33% SDS (50 mM Tris-HCl pH 8.0, 10 mM EDTA pH 8.0, 1.33% SDS). Crosslinks in eluted chromatin and input DNA were reversed by incubating at 65° overnight and treated with 8.4 μl of proteinase K (Merck) at 50° for 2 hr. DNA was purified using PCR purification kit (Qiagen) and further sonicated by Covaris to obtain DNA fragments averaging 150 bp. Multiplexed DNA libraries were prepared with NEBNext ChIP-seq library master mix prep set for Illumina (NEB) and NEBNext multiplex oligos for Illumina (NEB). ChIP and input DNA were sequenced by Illumina Miseq (4 samples per run, 50 nt single-end) using MiSeq reagent kit v2.

Single-end 50 nt reads were filtered and end-trimmed, followed by removal of the reads containing tag sequences, then mapped to the *S. cerevisiae* reference genome (sacCer2) as described (Ito *et al.* 2014). DNA enrichment (ChIP/Input) was calculated in 10-bp bins with normalization as described (Ito *et al.* 2014). A total of 1717 Red1 peaks were called as described (Murakami and Keeney 2014) using a 2010-bp Parzen (triangular) sliding window and a threshold of 1× genomic mean coverage.

### Data availability

Plasmids and strains are available on request. ChIP-seq data are available at the Gene Expression Omnibus (GEO) under accession numbers GSE100564 (H3 and H3 T11ph) and GSE103823 (Red1).

## RESULTS

### H3 T11 phosphorylation during meiosis is a response to DSBs

As part of a larger effort to identify meiotically regulated histone modifications in *S. cerevisiae*, we performed western blots on meiotic whole-cell extracts with antibodies to H3 T11ph. Under these conditions, signal was undetectable in mitotically cycling, premeiotic (G1-arrested, 0 hr), or early meiotic (through 2 hr) cultures, but accumulated transiently during meiosis with a maximum at ∼3 to 5 h (**Figure 1Ai, 1B**). This signal diminished as cells completed the first meiotic division (∼7 hr; **Figure 1Ai, 1B, 1C**). These findings agreed with studies reported while this work was in progress (Govin *et al.* 2010).

The anti-H3 T11ph signal occurred when DSBs are usually maximal under these conditions [∼3 to 5 hr (e.g., Thacker *et al.* 2014)], and coincided with an increase in H2A S129 phosphorylation (γ-H2A) (**Figure 1Ai**), which is formed by Mec1 and Tel1 kinases in response to meiotic DSBs (Mahadevaiah *et al.* 2001; Shroff *et al.* 2004). These results suggested H3 T11ph might be a DSB response, but H3 T11ph signal also coincided with an increase in H3 S10 phosphorylation (**Figure 1Ai**), which is DSB-independent (Hsu *et al.* 2000).

We therefore examined genetic requirements for H3 T11ph. The modification was undetectable in a strain with catalytically inactive Spo11 (*spo11-Y135F*; **Figure 1Aii, 1B**). As expected, induction of higher γ-H2A signal was not seen in *spo11-Y135F,* but H3 S10ph was induced (**Figure 1Aii**). H3 T11ph appeared in a *rad50S* strain, in which DSBs form but persist with unresected 5′ ends, so DSB resection is dispensable (**Figure 1Aiii**). H3 S10ph was unaffected in this mutant, but elevated γ-H2A levels persisted to late time points consistent with unmitigated Tel1 activity (Usui *et al.* 2001).

H3 T11ph appeared and disappeared in *rad51*Δ with kinetics similar to wild type (**Figure 1Av**), but persisted at high levels in *dmc1*Δ(**Figure 1Avi**) (a different antibody was used for these blots, discussed below). Both *rad51*Δ and *dmc1*Δ have defects in meiotic DSB repair (note the persistent γ-H2A), but with a more complete block in *dmc1*Δ (Bishop *et al.* 1992; Shinohara *et al.* 1992). Meiotic arrest is also nearly complete in *dmc1*Δ, whereas divisions occur in *rad51*Δ after a delay (**Figure 1C**) (Bishop *et al.* 1992; Shinohara *et al.* 1992).

To determine whether this persistent H3 T11ph signal was due to persistent DSBs or to meiotic arrest, we examined an *ndt80*Δ mutant. Ndt80 is a transcription factor needed for pachytene exit (Xu *et al.* 1995; Chu and Herskowitz 1998), and DSB repair defects cause arrest via checkpoint kinase-mediated inhibition of Ndt80 (Tung *et al.* 2000; Gasior *et al.* 2001). H3 T11ph did not persist at high levels in an *ndt80*Δ mutant; instead it peaked at 4 h at a slightly lower level than in wild type, then diminished to low residual levels similar to those seen at late times in wild type (**Figure 1Aiv**). This agrees with a recent report demonstrating H3 T11ph appearance and disappearance by western blotting and immunofluorescence of spread chromosomes (Subramanian *et al.* 2016). Therefore, high-level H3 T11ph persistence correlates with continued presence of meiotic DSBs (as in *dmc1*Δ), but not with arrest. This behavior contrasts with that of a different Mek1 substrate, Hed1, which remains phosphorylated in *ndt80*Δ mutants (Prugar *et al.* 2017). Both γ-H2A and H3 S10ph persisted at high levels in *ndt80*Δ (**Figure 1Aiv**) (Hsu *et al.* 2000), suggesting these modifications require pachytene exit for removal (Subramanian *et al.* 2016).

Because the H3 N-terminal tail has many potential modification sites (**Figure 1D**) and a different antibody not used in our studies cross-reacts between H3 T11ph, H3 S10ph and other modifications (Nady *et al.* 2008), we sought to validate the antibody specificity for the anti-H3 T11ph antibodies we used. Both the monoclonal and polyclonal anti-H3 T11ph antibodies were specific but did not detect H3 T11ph if S10 was also phosphorylated (**Supplemental Figure S1** and Materials and Methods). The monoclonal gave a more robust signal with less background for non-histone proteins (**Figure 1B**), so we used this antibody for most subsequent experiments. Two different anti-H3 S10ph antibodies recognized their cognate modification, but not if T11 was also phosphorylated. These anti-H3 S10ph antibodies showed significant cross-reactivity to other histones and modifications (**Supplemental Figure S1** and Materials and Methods).

To test if DNA lesions could also give rise to elevated T11 phosphorylation during vegetative growth, cells were treated with X-rays or camptothecin. These DNA damaging agents failed to yield a detectable level of H3 T11ph despite inducing DNA damage responses as evidenced by increased γ-H2A (**Figure 1E**). Thus, high levels of H3 T11ph in response to DNA damage are specific to meiosis. The strength of the meiotic H3 T11ph signal as compared to the undetectable levels under these blotting conditions for cycling or premeiotic cells or the *spo11-Y135F* mutant indicates that the amount of H3 T11ph formed in meiosis is vastly greater than what has been reported to be formed by pyruvate kinase during vegetative growth (Li *et al.* 2015).

### H3 T11ph in response to DSBs in *S. pombe* meiosis

To determine if meiotic H3 T11ph is evolutionarily conserved, we analyzed synchronous meiosis in *S. pombe* haploid *pat1-114* mutants (Bahler *et al.* 1991). H3 T11ph appeared transiently at ∼4– 5 hr after the initiation of meiosis and was not detected in a mutant lacking Rec12 (the Spo11 ortholog) or in vegetative growth (**Figure 2**). H3 T11ph appeared after a Rec12-dependent increase in γ-H2A that started around 3–3.5 hr, when DSBs typically appear under these conditions (Cervantes *et al.* 2000). (The initial wave of γ-H2A signal at or before 2 hr is Rec12-independent (**Figure 2B**) and possibly associated with DNA replication.) These results indicate that H3 T11ph forms in response to DSBs in *S. pombe*. H3 T11ph appeared and disappeared with apparently normal kinetics in a *rad50S* mutant in contrast to γ-H2A, which persisted at high levels (**Figure 2C**).

**Figure 2.**
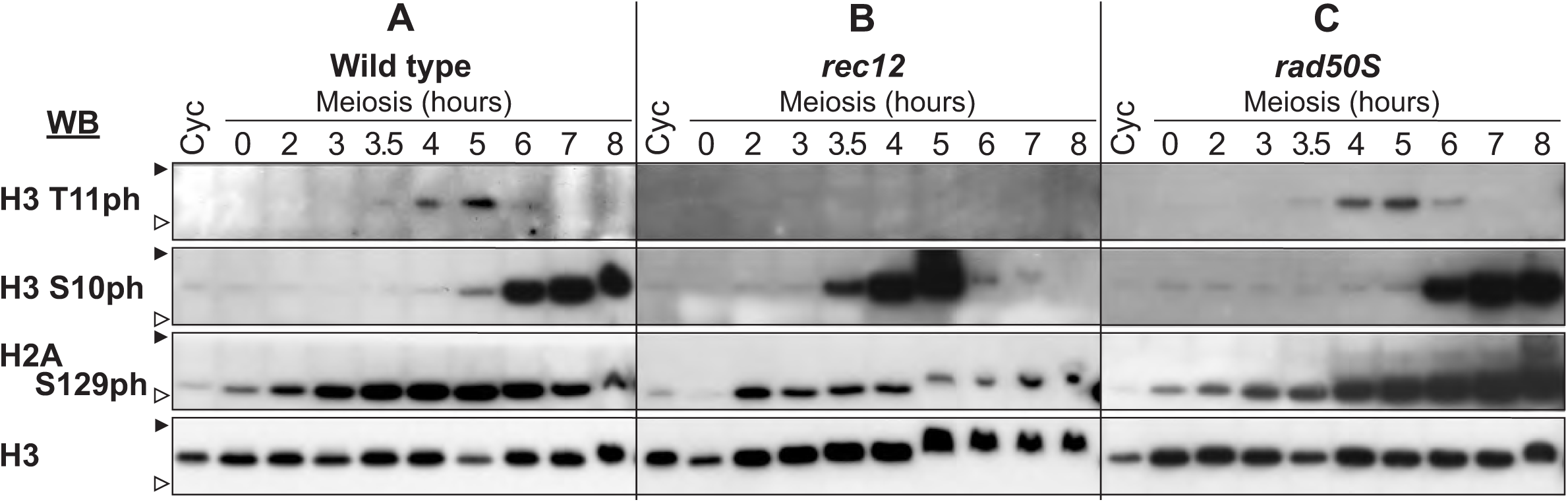
H3 T11 phosphorylation in *S. pombe* meiosis. Western blots of whole-cell extracts from haploid *pat1-114* strains undergoing synchronized meiosis. Antibodies used were the same as in **Figure 1Av**. Filled and open arrowheads indicate 20 and 15 kDa molecular weight markers, respectively. The altered electrophoretic mobility of histones at later time points in some cultures was probably caused by varying levels of contaminating DNA in the extracts rather than differential post-translational modifications.

H3 S10ph also appeared during meiosis, but unlike in *S. cerevisiae*, this modification occurred later than H3 T11ph (**Figure 2A**). In the *rec12* mutant, H3 S10ph was observed earlier than normal and was largely gone by 6 hr (**Figure 2B**). This result is consistent with accelerated meiotic progression in *rec12* mutants (Doll *et al.* 2008), and indicates that both appearance and disappearance of H3 S10ph are developmentally regulated.

### H3 T11 is a direct target of Mek1 kinase

The timing and genetic control of H3 T11ph in *S. cerevisiae* suggested that a meiosis-specific, DSB-responsive kinase was responsible. Mek1 expression coincides with H3 T11ph from 3–7 hr in meiosis (Carballo *et al.* 2008), and the T11 sequence context matches the Mek1 target consensus (RXXT; **Figure 1D**) (Mok *et al.* 2010; Suhandynata *et al.* 2016). We therefore treated a *dmc1*Δ strain expressing an ATP-analog sensitive *mek1* allele (*mek1-as*) with an inhibitor specific for the mutated Mek1 kinase, 1-NA-PP1 (Wan *et al.* 2004). Inhibitor addition at 4 hr caused rapid disappearance of H3 T11ph within the first hour (**Figure 3A**). This result demonstrates that Mek1 activity is necessary to maintain H3 T11 phosphorylation, and further implies that this modification is dynamic with a half-life much shorter than one hour.

**Figure 3.**
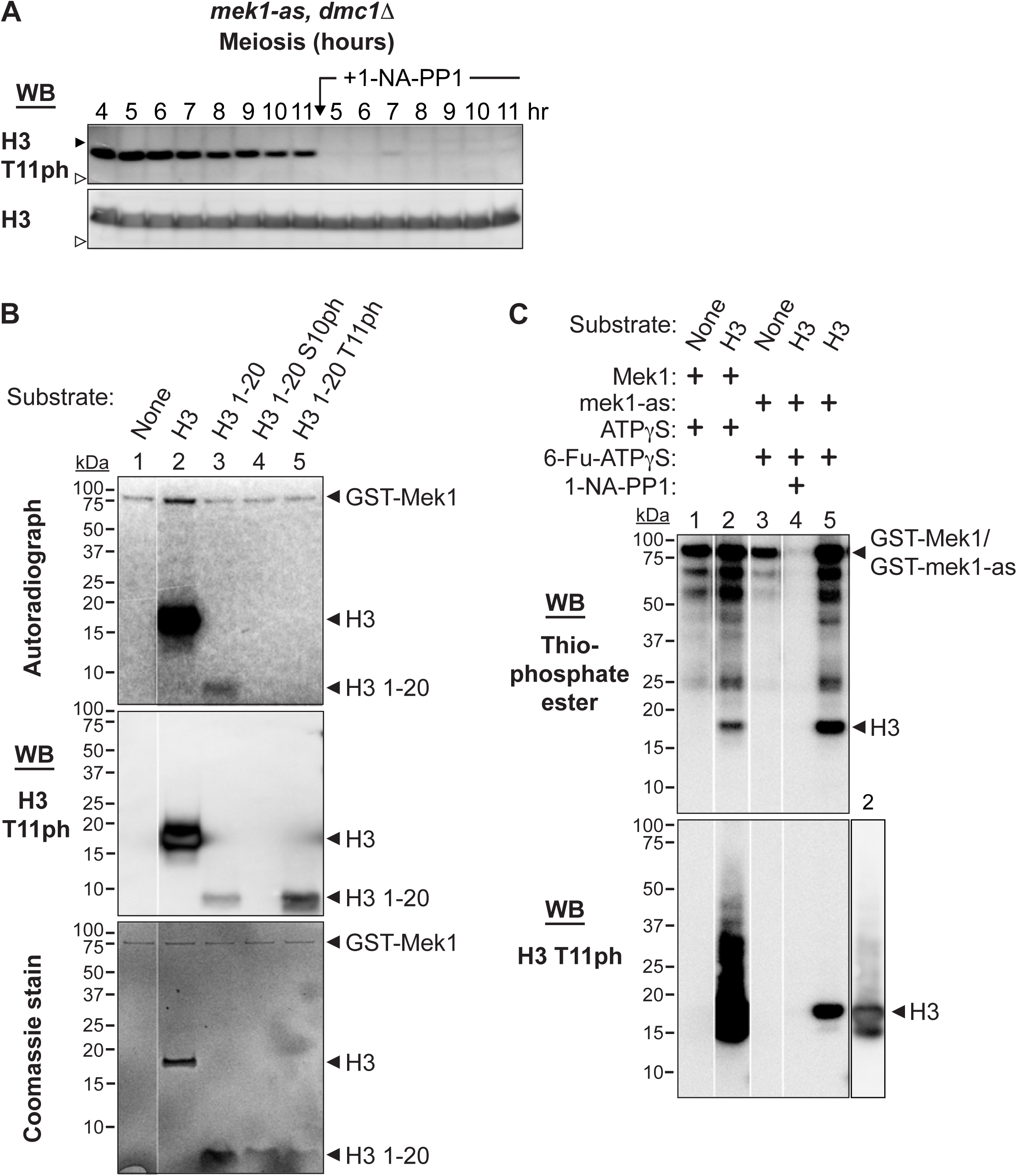
H3 T11 is a direct target of Mek1 kinase. **(A)** Persistence of H3 T11ph requires maintenance of Mek1 kinase activity. A meiotic culture of a *mek1-as*, *dmc1*Δ (strain SKY3095) was split 4 hr after transfer to sporulation medium. One part was left to continue in meiosis untreated, the other part was treated with 1 μM 1-NA-PP1. Whole-cell extracts were prepared at the indicated times and assayed for H3 T11ph by western blotting (mAb; EMD Millipore 05-789). Filled and open arrowheads indicate 20 and 15 kDa molecular weight markers, respectively. Numbers indicate hours after transfer to sporulation medium. **(B)** Mek1 kinase assay using radioactive ATP. Affinity-purified GST-Mek1 (250 ng) was incubated in the presence of [γ-^32^P]ATP either alone or with 2 μg recombinant H3 or 5 μg of unphosphorylated or phosphorylated synthetic H3 1-20 peptides as substrates. Reactions were separated by SDS-PAGE and visualized by autoradiography (top), anti-H3 T11ph western blot (middle; polyclonal Active Motif 39151), and Coomassie staining. **(C)** Mek1 kinase assay by semisynthetic epitope labeling. Kinase reactions were carried out with affinity-purified GSTMek1 (2 μg) or GST-Mek1-as (0.76 μg) in the presence of ATPγS or 6-Fu-ATPγS with 2 μg recombinant H3. After incubation 30 min at 30°C, PNBM (*p*-nitrobenzyl mesylate) was added to alkylate the thiophosphorylated target sites. Reactions were then separated by SDS-PAGE and analyzed by western blotting with anti-thiophosphate ester monoclonal antibody (top panel; Epitomics 2686-1). Because ATP was also included in all reactions, samples were also analyzed by western blotting with anti-H3 T11ph monoclonal antibody (EMD Millipore 05-789) to detect the subset of H3 proteins modified on T11 with phosphate instead of thiophosphate. Note that 1-NA-PP1 inhibits ability of GST-Mek1-as1 to use either ATP or Fu-ATPγS to phosphorylate H3 T11 (lane 4). A shorter exposure of lane 2 of the anti-H3 T11ph blot is shown to the right (asterisk indicates an H3 fragment present in the recombinant histone sample). Interstitial lanes were removed from images in panels B and C as indicated by the white lines.

This result agreed with prior findings demonstrating that H3 T11ph is reduced or absent in a *mek1*Δ mutant (Govin *et al.* 2010). However, these findings did not establish whether H3 T11 is a direct target of Mek1. To address this question, we carried out two types of *in vitro* kinase assay using GST-tagged Mek1 purified from meiotic *S. cerevisiae* cells (Wan *et al.* 2004; Niu *et al.* 2007). First, we used [γ-^32^P]ATP and full-length H3 or synthetic H3 peptides as substrates (**Figure 3B**). GST-Mek1 was visible in all lanes by Coomassie staining (**Figure 3B**, bottom panel) and its activity was confirmed by its ability to autophosphorylate **(Figure 3B**, top panel) (Niu *et al.* 2009). GST-Mek1 was able to phosphorylate full-length H3 and a peptide representing H3 amino acids 1-20 (**Figure 3B**, top panel, lanes 2 and 3). Phospho-transfer was specific for T11, as shown by western blot (**Figure 3B**, middle panel, lanes 2 and 3) and inability to label an H3 1-20 peptide that was already phosphorylated on T11 (**Figure 3B**, lane 5). Interestingly, GST-Mek1 was also unable to phosphorylate a peptide carrying a phosphate on S10 (**Figure 3B**, top panel, lane 4).

The second assay used a semisynthetic epitope system (Allen *et al.* 2007) to detect phosphorylation of H3 by Mek1. GST-Mek1 or GST-Mek1-as was incubated with recombinant H3 and the ATPγS analog, 6-Fu-ATPγS. Thiophosphates transferred by Mek1 to substrates were then alkylated to create an epitope that was detected on western blots with an anti-thiophosphate ester antibody (Niu *et al.* 2009; Lo and Hollingsworth 2011). Both GST-Mek1 and GST-Mek1-as exhibited autophosphorylation and phosphorylation of H3 (**Figure 3C**, lanes 2 and 5). Moreover, 1-NA-PP1 inhibited both autophosphorylation and H3 phosphorylation by GST-Mek1-as (**Figure 3C**, lane 4), ruling out the possibility of a contaminating kinase phosphorylating H3 T11. We conclude that H3 T11 is a direct substrate of Mek1.

### Limitations of a plasmid shuffle system for examining histone mutants

To determine the function of H3 T11 phosphorylation, we constructed strains carrying targeted mutations of T11 alone and in combination with other histone mutations. We initially tested an existing plasmid shuffle system (Ahn *et al.* 2005) by porting it to the SK1 strain background. In this approach, also used independently by others (Govin *et al.* 2010), the endogenous histone genes were deleted and complemented by wild-type histone genes on a *URA3*-marked *ARS-CEN* plasmid. Histone mutants were introduced on a separate *LEU2 ARSCEN* plasmid and loss of the *URA3* plasmid was selected for on medium containing 5-FOA.

However, this approach was sub-optimal because of the poor stability of the *ARS-CEN* plasmids in SK1. For example, when liquid cultures of the base histone-deletion strain carrying the *URA3* covering plasmid were grown under conditions selective for the plasmid (i.e., synthetic complete medium lacking uracil), plating on solid medium yielded an efficiency of only 67.2% ± 4.9% (mean ± SD of 5 replicates; colony-forming units per cell plated). Assuming that most cells that failed to form a colony were those that had lost the plasmid because of missegregation during mitosis, it is likely that plasmid copy number per cell is highly variable in the population. Cells with one vs. two copies of an H3/H4-encoding plasmid would likely differ in total histone protein levels and/or have different imbalances with endogenous H2A/H2B. Altered histone gene dosage can cause deleterious effects (Meeks-Wagner and Hartwell 1986; Clark-Adams *et al.* 1988), so it is possible that cell-to-cell heterogeneity in histone gene copy number might mask or exacerbate the effects of histone point mutations. Furthermore, differences in copy number might have a substantial effect on variation in viability of spores (see below). Finally, although cells in the culture that have lost the histone plasmid would be inviable and therefore presumably would not sporulate, they would contribute to population average measurements in physical assays of recombination.

To circumvent these limitations, we turned to mutagenesis methods that use gene replacement or stable chromosomal integration (Materials and Methods). Stable integration is relatively rapid and obviates concerns about plasmid stability and heterogeneous gene dosage, but may not fully recapitulate expression from endogenous histone gene loci. The gene replacement strategy provides an even cleaner manipulation of histone genotype, but is more cumbersome because it requires separately mutating two histone gene loci.

### Absence of H3 T11 phosphorylation causes little or no overt phenotypes by itself

We replaced both endogenous *H3* genes (*HHT1* and *HHT2*) with *hht1-S10A, T11V* and *hht2-S10A, T11V* mutant alleles to eliminate phosphorylation of both S10 and T11. This mutant expressed normal H3 protein levels and neither H3 S10ph nor H3 T11ph could be detected, as expected (**Figure 4A, lanes 3–4**). The mutant displayed normal vegetative growth (**Figure 4B**), similar to a recent report (Li *et al.* 2015). Surprisingly, however, the mutant also displayed normal spore viability (**Table 1**). Meiotic DSBs appeared in normal numbers and locations and disappeared with normal kinetics as assessed by Southern blotting of pulsed-field gels probed for chromosome III (**Figure 4C**), and meiotic progression was not delayed (**Figure 4D**). These results indicate that most if not all meiotic events occur efficiently in the complete absence of both S10ph and T11ph.

**Table 1.** Absence of H3 T11ph does not compromise spore viability.

| Mutation method <sup>a</sup> | H3 genotype <sup>b</sup> | Strain | Spore viability (no. of tetrads) |
| --- | --- | --- | --- |
| Replacement | Wild type | 2701 | 97% (44) |
|  | *S10A T11V* | 2705 | 97% (44) |
| *H3-H4* integration | Wild type | 3311 | 96% (22) |
|  | *S10A* | 3333 | 97% (64) |
|  | *T11V* | 3342 | 95% (64) |
|  | *S10A T11V* | 3334 | 93% (64) |
|  | *T11A* | 3312 | 95% (86) |
|  | *T11S* | 3313 | 99% (22) |
|  | *T11D* | 3332 | 97% (64) |
|  | *T11E* | 3303 | 97% (64) |
| Four-core integration | Wild type | 3330 | 94% (22) |
|  | *T11V* | 3264 | 97% (64) |
| | $\Delta N$ | 2388 | 81% (44) |
| Plasmid shuffle, Expt. 1 <sup>c</sup> | Wild type | 3438–3440 | 90% (90) |
|  | *T11A* | 3441–3443 | 86% (90) |
| Plasmid shuffle, Expt. 2 <sup>c</sup> | Wild type | 3438–3440 | 75% (92) |
|  | *T11A* | 3441–3443 | 77% (94) |
| Plasmid shuffle, Expt. 3 <sup>c</sup> | Wild type | 3438–3440 | 74% (96) |
|  | *T11A* | 3441–3443 | 76% (96) |

<sup>a</sup> Replacement: Both *HHT1* and *HHT2* were replaced with wild-type or mutant copies at their endogenous locations. Integration: Stable integration at *leu2::hisG* of a cassette carrying either the *H3* and *H4* gene pair *HHT2-HHF2*, or all four core histone genes *HTA1-HTB1* and *HHT2-HHF2*. Endogenous loci (encoding H3 and H4 or all four core histones, respectively) were deleted. Plasmid shuffle: Replacement of a *URA3* plasmid carrying wild-type *HHT2-HHF2* with a *LEU2* plasmid carrying either wild-type or mutant versions. The endogenous *H3* and *H4* loci were deleted.
<sup>b</sup> Genotypes are homozygous unless plasmid-based. See **Supplemental Table S1** for complete genotypes.
<sup>c</sup> Three independent 5-FOA<sup>R</sup> colonies were isolated for each shuffle plasmid and were dissected separately. The dissections were performed on three separate occasions by two different investigators; all six strains were dissected in parallel in each experiment. A breakdown of results by strain and experiment is provided in **Figure 4F**. Neither *H3* genotype nor clone identity was a significant predictor of altered spore viability ( $p \geq 0.9$ , linear regression).

**Figure 4.**
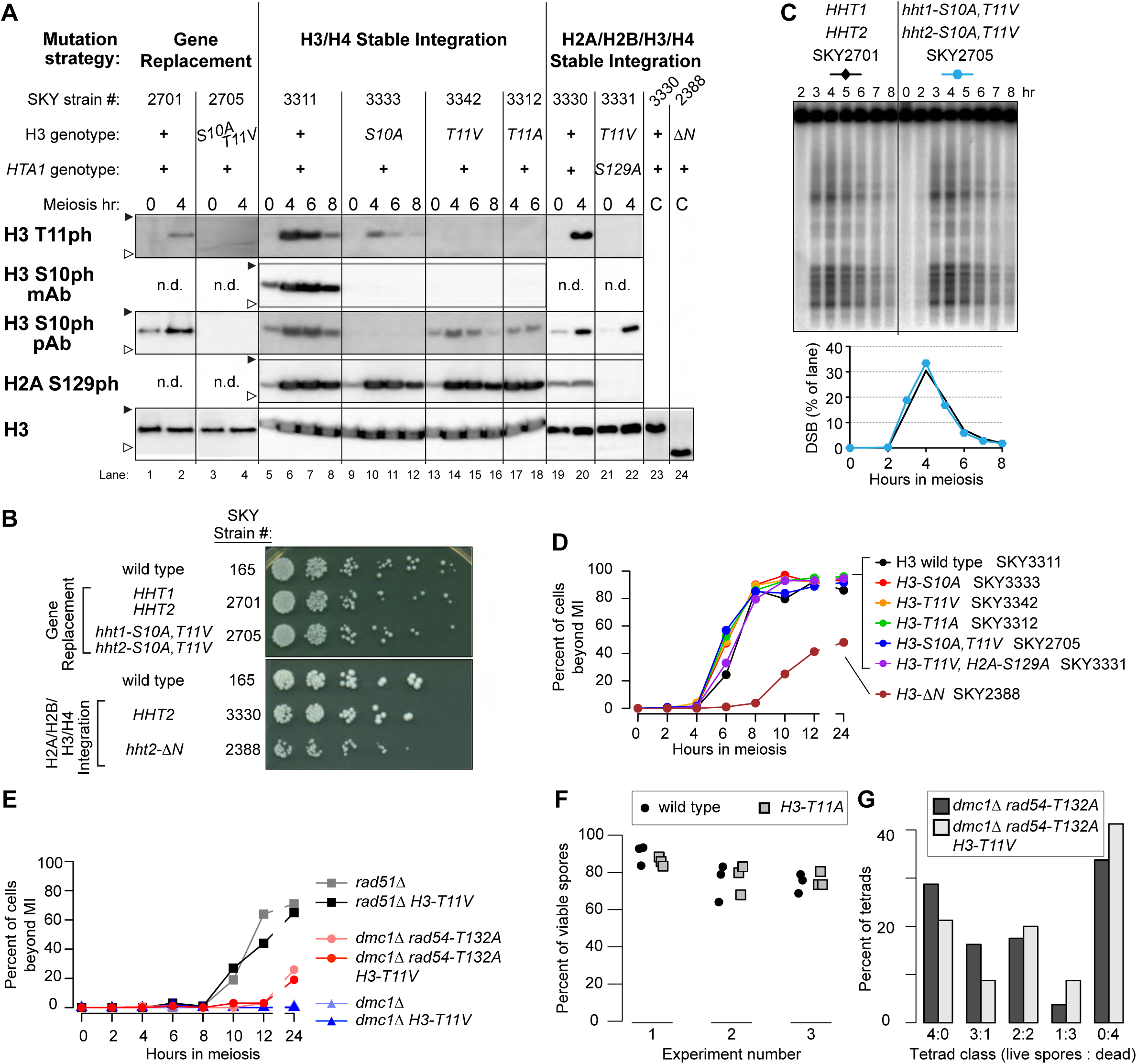
Characterization of histone mutant strains. **(A)** Composite of western blots of whole-cell extracts from synchronous meiotic cultures or asynchronous cycling vegetative cultures (“C”) carrying the indicated histone mutations. Antibodies used were: anti-H3 T11ph polyclonal (Active Motif 39151) or anti-H3 T11ph monoclonal (EMD Millipore 05-789); anti-H3 S10ph monoclonal (EMD Millipore 05-817); anti-H3 S10ph polyclonal (EMD Millipore 06-560); anti-γ-H2A (Abcam 15083); and anti-H3 (Abcam 1791). Filled and open arrowheads indicate 20 and 15 kDa molecular weight markers, respectively. “n.d.” indicates not determined; “*hht2-ΔN*” encodes H3 lacking its N-terminal 30 amino acids. **(B)** Vegetative growth of *H3* mutant strains. Cells from overnight cultures were spotted onto YPD plates using a manifold pin replicator and represent 1:5 serial dilutions starting with ∼2.5 × 10^6^ cells/ml. **(C)** Analysis of meiotic DSB formation. High-molecular-weight DNA isolated in agarose plugs was separated by pulsed-field gel electrophoresis followed by Southern blotting and indirect end-labeling with a probe directed against *CHA1* on the left arm of chromosome III. The lower panel shows quantification of the DSB signal as percent of lane total after background subtraction. (**D,E**) Meiotic progression of representative histone mutant strains. Cells were fixed and stained with DAPI and the fraction of cells with ≥ 2 nuclei was counted (n ≥ 100 cells per time point). For panel D, points represent mean of two independent experiments. For panel E, strains used were *rad51*Δ (SKY6218); *rad51Δ H3-T11V* (SKY6224); *dmc1Δ rad54-T132A* (SKY6210); *dmc1Δ rad54-T132A H3-T11V* (SKY3659); *dmc1*Δ (SKY6200); and *dmc1Δ H3-T11V* (SKY6220). **(F)** Spore viabilities in plasmid shuffle strains expressing wild-type H3 or H3 T11A. Three independent clones isolated for each genotype were sporulated and tetrads were dissected in three separate experiments. Each point represents the value from a single isolate (n = 30–32 tetrads per data point). See **Table 1** for summary and text for statistical test. Strains used were: *H3* wild type (SKY3438-3440) and *H3 T11A* (SKY3441-3443). **(G)** Evidence that the *H3-T11V* mutation increases MI nondisjunction in a *rad54-T132A dmc1Δ* background. The distribution of viable spores in tetrads is shown for the indicated strains. An increase in 2- and 0-spore-viable tetrads (rather than 3- or 1-spore-viable) suggests an increased frequency of MI nondisjunction. Strains were *dmc1Δ rad54-T132A* (SKY6210) and *dmc1Δ rad54-T132A H3-T11V* (SKY3659).

To more easily manipulate histone mutants, we used a chromosomal integration strategy to introduce genes for just *H3* and *H4* as a pair (*HHT2-HHF2*) or all four core histones (*HTA1-HTB1, HHT2-HHF2*) in strains deleted for the endogenous genes for *H3-H4* or all four histones. Wild-type or mutant histone genes were integrated on chromosome III at *LEU2*. Strains expressing H3-S10A, -T11V, or -T11A single mutant proteins or the H3-S10A T11V double mutant were examined in meiotic time courses for H3 S10 and T11 phosphorylation (**Figure 4A**). Importantly, H3 T11 could still be phosphorylated when S10 was mutated to alanine (**Figure 4A, lanes 9–12**); the lower signal in the anti-H3 T11ph western blot could reflect reduced T11 phosphorylation or decreased antibody affinity due to the changed epitope. Similarly, mutation of H3 T11 to alanine or valine did not prevent phosphorylation of S10, as detected with the polyclonal anti-H3 S10ph antibody, although recognition by the monoclonal anti-H3 S10ph antibody was sensitive to these mutations (**Figure 4A, lanes 13–18 and 21–22**).

As with gene replacement, all of these mutants yielded spore viabilities indistinguishable from matched wild-type controls (**Table 1**). Meiotic divisions were not delayed; if anything, divisions were slightly earlier in the *H3* mutants than the control strain (**Figure 4D**). Whether this difference has functional significance is unclear, especially given that *H3-T11* mutations did not alleviate arrest caused by DSB repair defects (see below). *H3-T11A* also supported wild-type interhomolog recombination between *arg4* heteroalleles [23 ± 1.5 Arg^+^ recombinants per 1000 viable cells for wild type (SKY3428) vs. 24 ± 0.8 for *H3-T11A* (SKY3431), mean ± SD for three independent cultures]. Other mutations of H3 T11 yielded similar results: changing T11 to serine or potential phosphomimetic residues (T11D or T11E) again yielded wild-type spore viability (**Table 1**). Mutating H3 T11 also did not reduce spore viability when combined with mutation of H2A S129 [which is also by itself largely dispensable for proper meiosis (Shroff *et al.* 2004; Harvey *et al.* 2005)] or with absence of the H3 K4 methyltransferase Set1 [which governs DSB distributions (Sollier *et al.* 2004; Borde *et al.* 2009; Acquaviva *et al.* 2013; Sommermeyer *et al.* 2013)] (**Figure 4A, lanes 21-22** and **Table 2**).

**Table 2.** Combining H3 T11 mutations with other mutations.

| **Additional mutation(s)<sup>a</sup>** | ***H3* genotype<sup>a</sup>** | **Strain** | **Spore viability**<br>**(no. of tetrads)** |
| --- | --- | --- | --- |
| *H2A S129A* | wild type | 3265 | 97% (64) |
|  | *T11V* | 3331 | 95% (64) |
| *set1Δ* | wild type | 4415 | 97% (42) |
|  | *S10A T11V* | 3329 | 97% (64) |
| *rad51Δ* | wild type | 3183 | 0% (44) |
|  | *S10A T11V* | 3186 | 0% (44) |
| *rad54-T132A dmc1Δ* | wild type | 6210 | 51% (80) |
|  | *T11V* | 3659 | 40% (80) |

<sup>a</sup> Genotypes are homozygous. See **Supplemental Table S1** for complete genotypes.

Mek1 is required for arrest or delay of meiotic progression when recombination is defective (Xu *et al.* 1997; Bailis and Roeder 2000). If H3 T11ph contributes substantially to this Mek1 function, then T11 mutations should alleviate some or all of the meiotic block in *rad51*Δ or *dmc1*Δ mutants. However, in cells lacking Rad51, the *H3-T11V* mutation had negligible effect on either the timing or efficiency of meiotic divisions (**Figure 4E**) and failed to rescue the spore inviability (**Table 2**). This H3 mutation also failed to alleviate the more stringent arrest in a *dmc1*Δ mutant (**Figure 4E**). Thus, H3 T11ph is dispensable for this checkpoint arrest function of Mek1.

Our findings differ from a prior report of an approximately 35% decrease in spore viability with plasmid-borne *H3-T11A* single or *S10A T11A* double mutants (Govin *et al.* 2010). We obtained the published *H3-T11A* plasmid and histone-deleted SK1 host strain (generously provided by J. Govin and S. Berger), verified the *H3-T11A* mutation by sequencing, and carried out the plasmid shuffle. Three independent 5-FOA-resistant clones for each genotype were sporulated and tetrads dissected for wild type and *H3-T11A* side-by-side. The experiment was repeated three times by two investigators. In our hands this *H3-T11A* mutant again yielded spore viability indistinguishable from the control with a wild-type *H3* plasmid (**Figure 4F and Table 1**, p > 0.9 by linear regression). However, unlike the normal spore viability observed in the stable integrant and gene replacement strains (**Table 1**), viability was consistently lower with plasmid-borne histone genes regardless of *H3* genotype (**Figure 4F and Table 1**). A similar baseline defect was reported previously (Govin *et al.* 2010). Furthermore, there was substantial heterogeneity in viability from experiment to experiment and between clones within each experiment (**Figure 4F and Table 1**). Within-experiment heterogeneity likely reflects stochastic culture-to-culture variability caused by plasmid instability. Between-experiment variability may reflect differences in sporulation conditions that in turn affect plasmid stability or the sensitivity of these strains to alterations in histone gene expression.

As a counter-example, we also examined a more extreme *H3* mutant in which the entire amino-terminal tail was deleted (*H3-ΔN*). The truncated histone was expressed at levels similar to full-length H3 in vegetative cells (**Figure 4A, lanes 23-24**). This mutant displayed vegetative growth defects (**Figure 4B**), delayed and less efficient meiotic divisions (**Figure 4D**), and reduced spore viability (**Table 1**; *p* = 0.0047, Fisher’s exact test).

### H3 T11ph contributes weakly to Mek1 function in the absence of Rad54 T132 phosphorylation

Because *H3-T11* mutations caused no overt defects on their own, we asked whether H3 T11ph might be redundant with Mek1 phosphorylation of Rad54 on T132 (Niu *et al.* 2009). A *rad54-T132A* mutation has little effect by itself, but in a *dmc1*Δ background it allows enough Rad51 activity to partially bypass arrest and produce some viable spores (Niu *et al.* 2009).

In a *rad54-T132A dmc1Δ* background, *H3-T11V* mutation significantly reduced spore viability (**Table 2**; *p* = 0.0087, Fisher’s exact test), with a decrease in four-spore-viable tetrads and an increase in two- and zero-spore-viable tetrads (**Figure 4G**). This segregation pattern suggests increased MI nondisjunction, as expected when intersister recombination is mediated by Rad51. In this context, *H3-T11V* gave no increase in overall meiotic division efficiency (**Figure 4E**).

These results suggest that H3 T11 phosphorylation provides a modest contribution to Mek1 function when meiotic recombination defects are encountered. Possible roles of H3 T11ph in these contexts are addressed in the Discussion. However, since the *H3-T11* mutation by itself does not detectably phenocopy a *mek1*Δ mutant, we conclude that H3 T11ph is normally dispensable for Mek1 function.

### H3 T11ph is enriched at axis-associated sites and, less strongly, along chromatin loops

H3 T11ph has been used as a cytological marker for Mek1 activity (Subramanian *et al.* 2016). Given our results establishing that H3 is a direct phosphorylation target of Mek1, we reasoned that H3 T11ph would also provide a sensitive and specific marker to reveal the genomic locations of active Mek1 kinase. We therefore assessed H3 T11ph genome-wide by ChIP-seq.

Samples were collected at 3 and 4 hr in meiosis from each of two independent wild-type cultures. To control for specificity of the H3 T11ph ChIP-seq signal, a sample was also collected from a 3.5-hr culture of a *spo11-Y135F* mutant. A set amount of *S. pombe* meiotic cells (4.5 hr in meiosis; 1% of the number of *S. cerevisiae* cells) was added to each *S. cerevisiae* cell sample prior to extract preparation. Mononucleosomes were liberated from formaldehyde-fixed meiotic chromatin by digestion with micrococcal nuclease (MNase), immunoprecipitated with the anti-H3 polyclonal or anti-H3 T11ph monoclonal antibodies, then the DNA was purified and deep sequenced and reads were mapped to the *S. cerevisiae* and *S. pombe* genomes. Each *S. cerevisiae* coverage map was normalized according to *S. pombe* read density for the same antigen from the same culture (**Figure 5A–C** and **Figure S2A,B**). The *S. pombe* spike-in control served two purposes. First, it helped minimize the effects of sample-to-sample variation in lysis, immunoprecipitation, and sequencing library preparation. Second, because the ratio of *S. cerevisiae* to *S. pombe* cells was fixed, the spike-in control provided a scaling factor to compare the relative yield of H3 or H3 T11ph between different *S. cerevisiae* samples (**Figure 5A–C**). Note that this allows comparison between samples for the same antigen, but does not quantify the yields of different antigens relative to one another. This approach was developed independently, but is similar to the previously described use of *Candida glabrata* as a spike-in control to calibrate *S. cerevisiae* ChIP-seq experiments (Hu *et al.* 2015).

**Figure 5.**
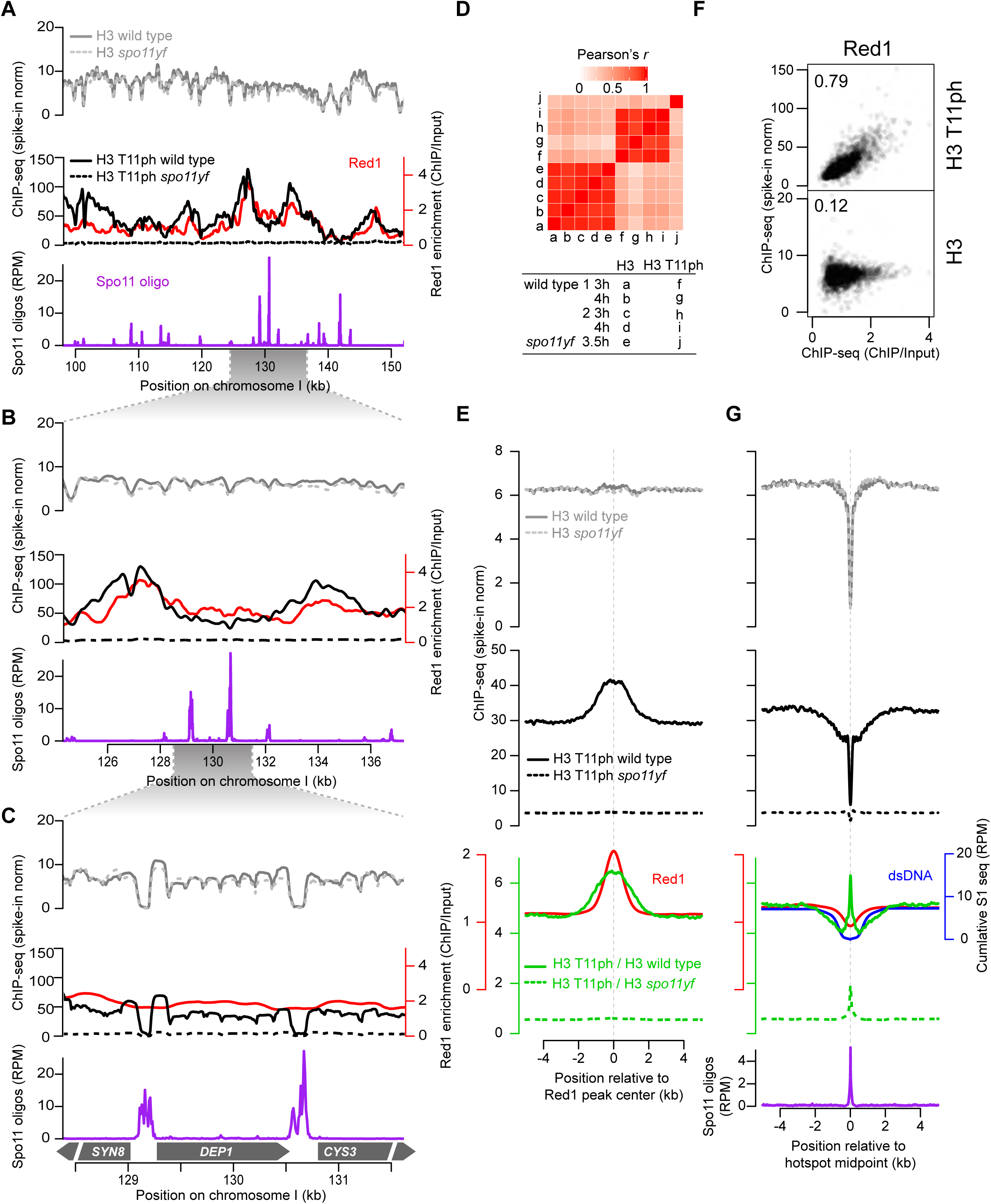
Spatial disposition of H3 T11ph along meiotic chromosomes. (**A–C**) Anti-H3 and anti-H3 T11ph ChIP-seq coverage across representative genomic regions. Coverage data for each chromosome were normalized (“norm”) relative to an *S. pombe* spike-in control. The four wild-type samples were averaged; the single *spo11-Y135F* sample (denoted “*spo11yf*”) is presented separately. For a given antigen (H3 or H3 T11ph), use of the internal *S. pombe* control allows direct quantitative comparison of relative yield in different samples, revealing in particular the DSB-dependent H3 T11ph signal via comparison of wild type with *spo11-Y135F*. The Spo11-oligo map (RPM, reads per million) (Mohibullah and Keeney 2016) and anti-Red1 ChIP-seq data (this study) are shown for comparison. The H3 and H3 T11ph data shown in A and B and all Red1 data were smoothed with 500 bp Parzen (triangular) sliding window. Color coding is retained in the other panels in this figure. (**D**) Reproducibility of histone ChIP-seq coverage maps. ChIP-seq coverage was averaged in 500 bp windows and compared between datasets. The heatmap is shaded according to the Pearson’s *r* value for each pairwise comparison. (**E**) H3 T11ph enrichment around presumed axis-attachment sites. H3 (upper graph) and H3 T11ph ChIP-seq coverage (middle graph) and smoothed (500-bp Parzen window) Red1 ChIP-seq data were averaged around 1717 Red1 ChIP-seq peaks. The green lines in the lower graph show the ratio of H3 T11ph to H3 ChIP-seq. (**F**) H3 T11ph correlates well with Red1 genome wide. Each point compares the H3 or H3 T11ph ChIP-seq coverage in wild type with Red1 ChIP-seq signal averaged across non-overlapping 5-kb bins. Correlation coefficients (Pearson’s *r*) are indicated in each plot. (**G**) H3 T11ph around DSB hotspots. ChIP-seq and Spo11-oligo data were averaged around Spo11-oligo hotspots and plotted as in panel E (Mohibullah and Keeney 2016); for clarity, hotspots more than 500 bp wide were excluded, and only the hottest 25% of hotspots are shown (N=872 hotspots). Note that vertical and horizontal scales for ChIP-seq data are the same in panels E and G to facilitate direct comparison. The blue line in the third panel shows the extent of dsDNA depletion predicted from S1-seq mapping of DSB resection tracts around the same hotspots (Mimitou *et al.* 2017). The lowest panel shows the average Spo11-oligo profile.

Several lines of evidence establish that these maps reported the distribution of H3 and DSB-dependent H3 T11ph with good specificity. At fine scale, H3 ChIP-seq coverage was low in promoters and showed prominent nucleosome-width peaks in coding sequences (**Figure 5C**), as expected for promoter-associated nucleosome-depleted regions (NDRs) and positioned nucleosomes in gene bodies (Jiang and Pugh 2009). Replicate samples agreed well, with all five H3 ChIP samples showing highly correlated distributions whether considered genome-wide (**Figure 5D**) or at individual loci (**Figure S2C**). For H3 T11ph, the four wild-type maps correlated well with one another but correlated poorly with either H3 ChIP-seq or H3 T11ph from *spo11-Y135F* (**Figure 5D**), as expected if the ChIP-seq signal was specific for this Mek1-dependent histone modification. Moreover, relative to the *S. pombe* spike-in, *S. cerevisiae* H3 ChIP-seq coverage was similar in wild-type and *spo11-Y135F* samples (**Figure S2C**), but H3 T11ph ChIP-seq coverage was substantially higher in all four wild-type samples than in *spo11-Y135F* (range of 3.4- to 11.6-fold across samples for genome-wide average) (**Figure 5A–C** and **Figure S2D**). The magnitude of the H3 T11ph signal (relative to spike-in) differed by up to ∼3.6 fold between the wild-type samples, possibly due to differences in read depth or in culture synchrony or efficiency. Nevertheless, the spatial patterns were highly reproducible (**Figure S2D**), so maps for wild type were averaged for further analysis.

At fine scale, H3 T11ph ChIP coverage showed depletion in NDRs and nucleosomal peaks at similar positions as in the H3 map (**Figure 5C**). This pattern is as expected since presence of a nucleosome (as revealed by bulk H3 localization) is a prerequisite for placement of H3 T11ph by Mek1. However, when maps were examined at larger size scales, H3 T11ph showed broad hills and valleys that were not matched in the H3 ChIP-seq (**Figure 5A,B**), revealing that H3 T11ph tends to be relatively enriched or depleted in domains several kb in width.

A priori, we envisioned two non-exclusive scenarios that might describe H3 T11ph localization: Enrichment at chromosome axes because that is where Mek1 protein is enriched cytologically and Mek1 interacts with axis proteins (Bailis and Roeder 1998; Wan *et al.* 2004; Carballo *et al.* 2008); or enrichment centered on DSB hotspots because Mek1 activation is a response to DSBs and Mek1 regulates DSB repair. We examined each possibility in turn.

#### Axis-associated sites

To test if H3 T11ph is enriched near axes, we compared its ChIPseq signal with the genome-wide distribution of axis component Red1. For this purpose, we used new ChIP-seq data acquired at 3 hr in meiosis from a strain carrying flag-tagged Red1 (**Fig 5A–C**) as well as earlier tiling microarray-based ChIP data (ChIP-chip) (Panizza *et al.* 2011). Spatial patterns for both ChIP methods agreed well (**Figure S3A,B**). The sites where ChIP signals for Red1 and other axis proteins are enriched are generally assumed to be the chromatin loop bases that are embedded in the chromosome axis (Blat *et al.* 2002; Panizza *et al.* 2011; Sun *et al.* 2015). These sites often but not always overlap with intergenic regions between convergent transcription units, presumably because transcription can push cohesin and associated axis proteins along chromosomes (Lengronne *et al.* 2004; Bausch *et al.* 2007; Sun *et al.* 2015).

Across individual chromosomal segments, peaks and valleys in the H3 T11ph signal appeared to correspond with peaks and valleys of Red1 (**Figure 5A**). Confirming this impression, average H3 T11ph signal formed a broad peak ∼4 kb wide when centered on Red1 ChIP-seq peaks, slightly wider than the average of Red1 itself (**Figure 5E**). No such enrichment was observed in the *spo11-Y135F* mutant (**Figure 5A,E**). H3 ChIP-seq showed no enrichment centered on Red1 peaks and was indistinguishable in wild type and *spo11-Y135F* (**Figure 5E**). Thus, H3 T11ph signal remained elevated at Red1 peak positions after correcting for bulk H3 levels (green line in **Figure 5E**). Furthermore, H3 T11ph ChIP-seq correlated well genome-wide with Red1 ChIP-seq, whereas H3 ChIP-seq correlated poorly with Red1 (**Figure 5F**). Similar results were obtained by comparing H3T11ph signal to ChIP-chip data for Red1 and another axis component, Hop1 (**Figure S3C,D**). We conclude that H3 T11ph is particularly prevalent where Red1 and Hop1 are enriched, and thus that Mek1 is highly active at axis-associated sites.

#### Around DSB hotspots

To test if H3 T11ph is enriched near DSB sites, we compared its ChIP-seq signal with DSB maps generated by sequencing of Spo11 oligos (Pan *et al.* 2011; Mohibullah and Keeney 2016). When centered on Spo11-oligo hotspots, histone ChIP-seq coverage showed a complex pattern of highly localized enrichment and depletion (**Figure 5G**). The average for total histone H3 showed strong depletion in hotspot centers, flanked by shallow alternating peaks and valleys (gray line in **Figure 5G**). This is the expected pattern from prior studies, reflecting the strong preference for DSBs in *S. cerevisiae* to form in promoter NDRs that are flanked by positioned nucleosomes (Ohta *et al.* 1994; Wu and Lichten 1994; Pan *et al.* 2011) (e.g., **Figure 5C**). [For clarity, the plots show averages for the hottest 25% of all hotspots after excluding unusually wide hotspots (> 500 bp); qualitatively similar results were obtained if all hotspots were averaged (data not shown).]

The average H3 T11ph ChIP-seq signal differed from this pattern in informative ways (black line in **Figure 5G**). First, at all positions across the averaging window, H3 T11ph ChIP-seq signal was much higher in wild type than in *spo11-Y135F*, and this difference was greater for stronger hotspots than for weaker ones (**Figure 5G**). Therefore, there is substantial DSB-dependent (thus presumably Mek1-dependent) H3 T11 phosphorylation all across the regions where DSBs usually form, not just at nearby axis sites.

Second, relative to the baseline genomic H3 T11ph signal, there was strong depletion at hotspot centers, indicated by the narrow cleft (∼200 bp wide) in the average profile (**Figure 5G**). This cleft corresponded well to the central cleft in the H3 ChIP-seq average, so we infer that this narrow zone of depletion reflects the fact that there are few histones available to be phosphorylated within the NDRs where hotspots generally occur. There was a peak at hotspot centers when the H3 T11ph levels were normalized to bulk H3 signal (green line in **Figure 5G**). However, there was also a peak when the *spo11-Y135F* map was normalized for bulk H3, thus much or all of this is a DSB-independent signal. This may be a ChIP-seq artifact, or could reflect a low level of Mek1-independent H3 T11ph enriched near promoters (Li *et al.* 2015). [Although gene promoters have lower nucleosome occupancy compared with the rest of the genome, they are not devoid of nucleosomes. For example, some promoters contain positioned, high-occupancy nucleosomes; some contain nucleosomes but only in a fraction of the population; and some contain sub-nucleosomal histone particles (Jiang and Pugh 2009; Floer *et al.* 2010; Weiner *et al.* 2010).]

Third, there was a broader zone of lower H3 T11ph signal flanking the central NDR and extending ∼2 kb on either side (**Figure 5G**). This zone extended into areas where bulk H3 levels were high, so the difference map (normalizing H3 T11ph to H3) revealed depletion for H3 T11ph relative to immediate surroundings (green line in **Figure 5G**). Nonetheless, the H3 T11ph signal across this region was substantially higher in wild type than in *spo11-Y135F*. Much of this depleted zone corresponded to the same areas degraded by exonucleolytic resection as measured by S1-seq (Mimitou *et al.* 2017) (blue line in **Figure 5G**). This suggests that some or all of this depletion reflects disruption of chromatin — and thus of ChIP-detectable H3 T11ph signal — accompanying DSB resection. Interestingly, the H3 T11ph depletion zone correlated with the dimensions of a zone that was also relatively depleted for Red1 at 3 hr in the ChIP-seq data (**Figure 5G**) and in the ChIP-chip data (**Figure S3E**).

Collectively, these findings suggest that the distribution of DSB-provoked H3 T11 phosphorylation is governed largely by the distribution of Red1 and other proteins that are directly involved in Mek1 activation. Further implications are addressed in the Discussion.

### H3 T11ph correlates with DSB frequency across large sub-chromosomal domains

We next examined larger scale variation in H3 T11ph ChIP signal across chromosomes. H3 T11ph ChIP signals were binned in non-overlapping windows of varying sizes from 0.5 to 40 kb, then compared (Pearson’s *r*) to Spo11-oligo densities or ChIP signals for Red1, Hop1, or Rec8 in the same bins (**Figure 6**). Comparisons using the ratio of H3 T11ph to H3 show which correlations are specific for the histone modification ChIP per se (green points in **Figure 6**) as opposed to underlying (background) enrichment or depletion in the bulk chromatin map (total H3; gray points). Comparisons using the ratio of wild type to *spo11-Y135F* for H3 T11ph show which correlations are specific for DSB-dependent (and thus Mek1-dependent) signal (blue points in **Figure 6**).

**Figure 6.**
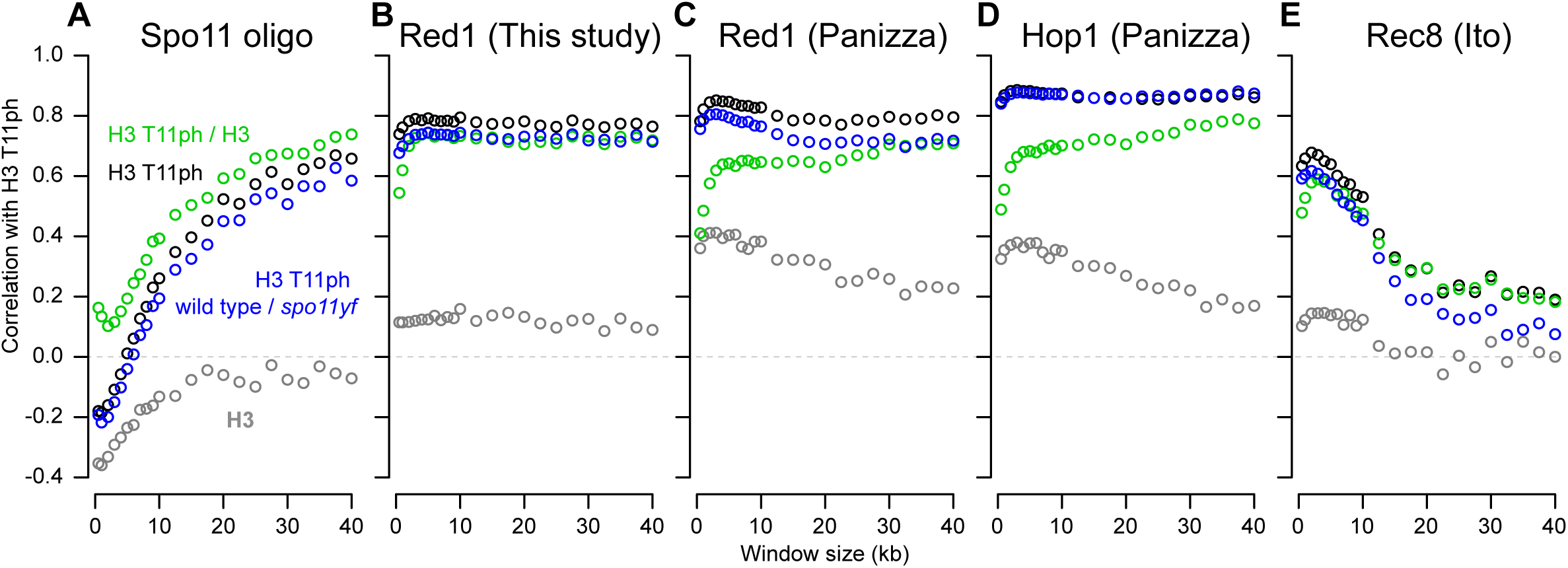
Scale-dependent correlations of H3 T11ph with chromosomal features. Anti-H3 (gray points) and anti-H3 T11ph (black points) ChIP-seq coverage was binned in non-overlapping windows of varying sizes and compared (Pearson’s *r*) to Spo11-oligo density (A) or ChIP-chip or ChIP-seq signals for Red1 (B, C), Hop1 (D), or Rec8 (E) averaged across the same windows. Red1 ChIP-seq data are from this study. Spo11-oligo (Mohibullah and Keeney 2016), Red1 and Hop1 ChIP-chip (Panizza *et al.* 2011), and Rec8 ChIP-seq data (Ito *et al.* 2014) were from previous studies. Green points show correlations using the ratio of H3 T11ph to H3 in wild type; blue points show correlations for the ratio of wild type to *spo11-Y135F* H3 T11ph signal.

For small windows (< 2 kb), both H3 and H3 T11ph were anticorrelated with Spo11-oligo density (**Figure 6A**). This pattern is driven by strong preference for DSBs to form in NDRs, and the attendant depletion of histone signal around hotspots (**Figure 5G**). In contrast, with large windows the H3 T11ph signal instead had a significant positive correlation with Spo11-oligo density, with Pearson’s *r* values high over a range of ∼25–40 kb (**Figure 6A**). This correlation was also high when the wild-type H3 T11ph ChIP data were normalized to coverage in *spo11-Y135F*, but no such correlation was seen for total histone H3, thus this pattern is specific for DSB-dependent H3 T11 phosphorylation. We infer that subchromosomal domains tens of kb wide that experience more DSBs also incur more Mek1 activity on average. This finding fits with the expectation that H3 T11ph is a faithful molecular reporter of DSB-provoked Mek1 kinase activity.

In contrast to the wide variation in correlation behavior depending on window size when H3 T11ph was compared to Spo11-oligo density, comparisons with either Red1 or Hop1 ChIP showed strong positive correlations over all window sizes tested (**Figure 6B–D**). Qualitatively similar results were obtained with Red1 ChIP-seq and ChIP-chip data, the principal difference being that the Red1 ChIP-chip data showed a higher but still weak correlation with bulk histone H3 ChIP for smaller window sizes (compare **Figures 6B,C**). Higher resolution and better specificity of sequencing vs. hybridization-based detection may explain this subtle difference between the ChIP-seq and ChIP-chip data for Red1. The good concordance of the ChIP-seq and ChIP-chip data for Red1 validates the use of published Hop1 ChIP-chip data. Hop1 was highly similar to Red1 in showing a largely scale-independent positive correlation with H3T11ph (**Figure 6C,D**).

Using Rec8 ChIP-seq data previously collected at 4 hr (Ito *et al.* 2014), H3 T11ph showed a positive correlation for short windows (less than ∼10 kb) but only weak correlation with larger windows (**Figure 6E**). This pattern can be understood as the combination of two spatial correlations with different length dependencies. At short distances (<10 kb), Mek1 activity is particularly enriched at preferred binding sites for Red1, Hop1, and Rec8 (i.e., presumptive axis attachment sites; **Figure 5E**). At longer distances (tens of kb), the domains that are relatively DSB-rich (and thus have more Mek1 activity) are also enriched for Red1 and Hop1 but not for Rec8 (Blat *et al.* 2002; Pan *et al.* 2011; Panizza *et al.* 2011; Ito *et al.* 2014).

## DISCUSSION

This study and others (Govin *et al.* 2010; Subramanian *et al.* 2016) establish that H3 T11 phosphorylation is highly induced during meiosis in *S. cerevisiae*. Our findings additionally demonstrate that H3 T11ph is a direct product of DSB-induced activation of Mek1. Mek1 is conserved in *S. pombe* (Perez-Hidalgo *et al.* 2003), so it seems likely that this kinase is also responsible for the H3 T11ph we observed in fission yeast.

Mek1 appears specifically in fungal taxa, but the larger Rad53 kinase family is ubiquitous in eukaryotes (Subramanian and Hochwagen 2014). Another member of this family, CHK1, was reported to be required for H3 T11ph in mouse fibroblasts (Shimada *et al.* 2008). In this case, however, DNA damage caused a decrease in H3 T11ph levels. It remains unknown if CHK1 directly phosphorylates H3 T11 or if H3 T11ph occurs in response to DSBs in mammalian meiosis. H3 T11ph has been reported during meiosis in sciarid flies (Escriba *et al.* 2011), suggesting evolutionary conservation beyond yeasts.

H3 T11 can also be directly phosphorylated by pyruvate kinase M2 in *S. cerevisiae* and mammalian cells, possibly to coordinate chromatin structure and gene expression with the cell’s nutritional status (Yang *et al.* 2012; Li *et al.* 2015). In cultured human cells, H3 T11ph is also formed by protein-kinase-C-related kinase 1 near promoters of androgen receptor-modulated genes (Metzger *et al.* 2008), and by death-associated protein (DAP)-like kinase during mitosis, particularly near centromeres (Preuss *et al.* 2003). Our results establish that meiotic induction of H3 T11ph in yeasts is fundamentally distinct from these other modes of H3 T11 phosphorylation in terms of provenance and genomic distribution.

### Possible functions of H3 T11ph in meiosis

Under the conditions in this study, histone mutations that eliminated H3 T11 phosphorylation caused no discernible meiotic defects by themselves. This was true with multiple independent mutagenesis strategies and numerous mutant constructs encoding different amino acid substitutions alone or in combination with mutation of H3 S10. We conclude that H3 T11ph is dispensable for meiosis under our standard conditions.

Why our results differed from a previous report (Govin *et al.* 2010) remains unknown. One possibility is that the highly variable spore viability in the plasmid shuffle system fortuitously gave the incorrect appearance of a meiotic defect in the earlier study. The reported decrease in spore viability [from ∼80% in the control to ∼50% with *H3-T11A* (Govin *et al.* 2010)] was of comparable magnitude to the intrinsic experimental variability we observed with plasmid-borne histone cassettes. Alternatively, studies in the two laboratories may have had undocumented differences in sporulation conditions to which *H3*-*T11* mutants are specifically sensitive.

Despite H3 T11ph being dispensable in unperturbed meiosis, we did observe that the *H3-T11V* mutation modestly exacerbated the phenotype of a *dmc1Δ rad54-T132A* mutant. One possibility is that H3-T11V protein acts as a weak competitive inhibitor of Mek1, thereby attenuating its ability to phosphorylate other substrates. However, we favor the alternative that the effect of the *H3-T11V* mutation is attributable to absence of H3 T11 phosphorylation per se. Possibly, H3 T11 helps Mek1 maintain residual interhomolog bias when Rad51 is the sole source of strand exchange activity. In this model, increased MI nondisjunction is caused by more of the residual DSB repair being between sister chromatids, and less between homologs. This interpretation is motivated by the increased intersister recombination observed in a *rad54-T132A* mutant when Mek1 activity is inhibited, and by the ability of the *rad54-T132A* mutation to rescue some spore viability in a *dmc1*Δ background but not in *dmc1Δ mek1Δ* (Niu *et al.* 2009). These findings indicate that other Mek1 targets contribute to interhomolog recombination by Rad51 when Dmc1 is missing and Rad54 cannot be phosphorylated. The recent discovery that Mek1 phosphorylates Hed1 and histone H2B make these strong candidates for additional redundancy (Callender *et al.* 2016; Suhandynata *et al.* 2016) (N.M.H., unpublished data).

If H3 T11ph does promote Mek1 function, albeit in a minor way, what might its role be? One possibility is that it is an effector of Mek1 signaling. This could be via recruitment to chromatin of proteins with phosphothreonine binding motifs such as the FHA domain, which is present in numerous proteins in *S. cerevisiae* including the recombination protein Xrs2 (Mahajan *et al.* 2008; Matsuzaki *et al.* 2008). Or, H3 T11ph might impinge on nucleosome stability, higher-order chromatin organization, or ability to install, remove, or read other histone modifications. We observed potential crosstalk between histone modifications in that H3 S10ph blocked the ability of Mek1 to phosphorylate T11 on the same peptide. Crosstalk of H3 T11ph with other H3 modifications has been documented in vegetatively growing yeast [H3 K4 methylation (Li *et al.* 2015)] and in human cells [H3 K9 acetylation (Yang *et al.* 2012) and demethylation (Metzger *et al.* 2008)]. A second, non-exclusive possibility is that H3 T11ph might maintain or amplify Mek1 activity via positive feedback. For example, the FHA domain of Mek1 might bind directly to H3 T11ph in a manner that stabilizes or increases the amount of active Mek1. Both general types of role — downstream effector or feedback amplifier — are compatible with the observed genetic interaction of *H3-T11* mutation with *dmc1Δ rad54-T132A*.

### Spatial organization of Mek1 activity

Although H3 T11 can apparently be phosphorylated by other kinases, the magnitude of the DSB- and Mek1-dependent signal combined with its rapid disappearance when Mek1 is shut off made H3 T11ph an excellent candidate for a molecular marker of ongoing Mek1 activity. Our experiments establish proof of principle for this use in genomic experiments, and also validate that cytological staining for H3 T11ph provides a direct readout of Mek1 activity (Subramanian *et al.* 2016).

The most prominent sites of H3 T11ph, and thus of Mek1 activity, were coincident with peaks of Red1 and Hop1, i.e., presumed axis attachment sites. This pattern is not surprising given that Mek1 protein appears to be enriched on axes as assessed by immunocytology (Bailis and Roeder 1998). However, immunolocalization does not reveal kinase activity per se, and cannot evaluate the degree to which activity might spread in *cis*. Interestingly, the H3 T11ph enrichment was similar to that of Red1 and Hop1 around axis sites. This more constrained pattern contrasts with the broad, relatively unstructured spread of γ-H2A over tens of kb around DSBs in yeast (Shroff *et al.* 2004). The local coincidence between enrichment patterns for H3 T11ph with those for Red1 and Hop1 could be because Mek1 protein is constrained, i.e., it rarely diffuses away from the sites where it has been activated. Alternatively, Mek1 may be rapidly inactivated if it diffuses away and/or the phosphates that Mek1 places outside the immediate vicinity of Mek1 activation sites might be more rapidly removed by phosphatases. In contrast, the broader γ-H2A distribution suggests that Mec1 and/or Tel1 activity can spread relatively uniformly around a DSB. We note that our analysis does not allow us to definitively evaluate how far H3 T11ph spreads around any given DSB, because we are looking at a population average across cells with DSBs at many different positions. It is conceivable that H3 T11ph (and thus Mek1 activity) spreads as far as Mec1/Tel1 activity does, given the strong correlation of H3 T11ph with Spo11-oligo density across large size scales. Rigorously addressing this question would require measuring H3 T11ph around a unitary, defined DSB.

We additionally observed substantial levels of DSB-dependent H3 T11ph across areas in between Red1 peaks, i.e., along the lengths of presumptive chromatin loops. This signal correlated with Spo11-oligo frequency (i.e., local DSB density) but was lower immediately around DSB hotspots, coincident with regions also depleted of Red1 in both the ChIP-seq and ChIP-chip datasets analyzed. For any protein, ChIP enrichment at given sites does not imply that these are the only sites of binding. In yeast meiosis, axis proteins Red1 and Hop1 are also detected above background across regions in between the sites of their principal enrichment. ChIP provides a population average measurement, so possible interpretations are that Red1 et al. are bound to different sites in different cells, but only at discrete sites in any one cell (an “axis sites only” model); or that there is lower level binding to chromatin loops in addition to higher level binding at axis sites (an “axis plus dispersed binding” model).

The DSB-dependent H3 T11ph ChIP signal tracked closely with Red1 and Hop1 ChIP at all size scales, both quantitatively and spatially. Thus, our findings are consistent with Mek1 activity manifesting anywhere Red1 and Hop1 are present to support it once DSBs have formed. It is interesting to note that the sites of highest Mek1 activity (i.e., Red1/Hop1 peaks) are spatially distinct from the DSB sites where Mek1 exerts its known biological function — Hop1/Red1-dependent control of recombination outcome. However, even though H3 T11ph ChIP signal was less abundant immediately adjacent to DSB sites than elsewhere, the DSB-proximal signal was substantially higher in wild type than in the *spo11-Y135F* control. Thus, we infer that active Mek1 kinase has access to chromatin and chromatin-associated proteins immediately surrounding DSBs.

A puzzle about the signal immediately adjacent to hotspots is that DSBs are exonucleolytically resected for ∼800 nucleotides on average on both sides of the break (Zakharyevich *et al.* 2010; Mimitou *et al.* 2017), but ssDNA should not be revealed in our ChIP-seq data even if it were still bound by histones because the sequencing library preparation protocol is not expected to support recovery of ssDNA. What then is the source of H3 T11-phosphorylated nucleosomes immediately around hotspots? Likely candidates are the sister of the broken chromatid, one or both intact chromatids of the homologous chromosome with which recombination is occurring, and/or recombination intermediates (D-loops and double Holliday junctions) if these are chromatinized. Because Mek1 controls homolog bias, we speculate that some or all of the H3 T11ph signal around hotspots is from Mek1 action on the sisters of broken chromatids. In support of this idea, we further note that the areas expected to be covered by DSB resection are also areas where the H3 T11ph signal is locally depleted. This pattern is thus in line spatially with what would be predicted if both sister chromatids are being exposed to Mek1 activity.

Our data are consistent with spreading of DSB-provoked Mek1 activity in *cis* along chromatin and concentrating wherever Red1 and Hop1 are also concentrated. The findings neither refute nor support the TLAC model (see Introduction), but are consistent with this model provided that DSB-dependent activation of Mek1 at axis sites can be accompanied by spreading of Mek1 activity to surrounding chromatin as well. Current versions of the TLAC model favor the idea that tethering occurs before DSB formation because some partners of Spo11 are enriched at axis sites rather than at hotspots but can be connected to hotspots physically via interactions with a reader (Spp1) of the H3 K4 methylation that is prominent around promoters (Panizza *et al.* 2011; Acquaviva *et al.* 2013; Sommermeyer *et al.* 2013). Such loop-axis interactions prior to DSB formation could provide a means to rapidly and specifically activate Mek1 at a nearby axis site in response to a DSB at a hotspot within a tethered loop.

Immunostaining experiments demonstrated H3 T11ph foci of variable size and intensity on chromatin where synaptonemal complex had not yet formed, although spatial disposition of this phosphorylation relative to axes has not been reported (Subramanian *et al.* 2016). Combining these cytological findings with our genomic data suggests that each DSB provokes a relatively large zone of Mek1 activation that encompasses the broken chromatin loop, its sister chromatid, and the adjacent loop bases(s). We speculate that this zone may extend across one or more loops, perhaps dependent in part on availability of sufficient amounts of Red1 and Hop1 to support Mek1 activity.

In summary, the detection of H3 T11ph is useful as an indicator of meiotic DSB formation, an indicator of Mek1 activation level, and a marker of the spatial organization of chromatin that Mek1 acts upon. H3 T11ph ChIP will be a powerful tool for dissecting not only the function of Mek1 but also the higher order structural organization of recombining chromosomes.

## ACKNOWLEDGMENTS

We thank J. Song (Keeney lab) for assistance with strain construction and ChIP-seq sample preparation; K. Kugou (Ohta lab) for assistance with strain construction and Red1 ChIP-seq data; A. Oda (Ohta lab) for assistance with Red1 ChIP-seq data; F. Klein (University of Vienna) for suggesting *S. pombe* as a spike-in control for ChIP-seq; G. Smith (Fred Hutchinson Cancer Research Center) for *S. pombe* strains and advice on culturing; J. Govin and S. Berger (University of Pennsylvania) for providing histone mutant plasmids and strains and for communicating information prior to publication; C. Hughes and C.D. Allis (Rockefeller University) for providing peptides, recombinant histones, plasmids and valuable guidance; N. Hunter (University of California, Davis) and G. Chung (Keeney lab) for strains; A. Viale and N. Mohibullah (MSKCC Integrated Genomics Operation) for sequencing; and S. Yamada (Keeney lab) and N. Socci (MSKCC Bioinformatics Core) for bioinformatics assistance. MSKCC core facilities were supported by NIH/NCI Cancer Center Support Grant P30 CA008748. R.K. was supported in part by NIGMS predoctoral training award T32 BM008539. This work was supported by NIH grants R01 GM058673 and R35 GM118092 (to S.K.) and R01 GM050717 (to N.M.H.), and by grants from the Japan Society for the Promotion of Science (15H04625, 26291018, 17H03711 to K.O.).

## SUPPLEMENTAL FIGURE LEGENDS

**Figure S1.**
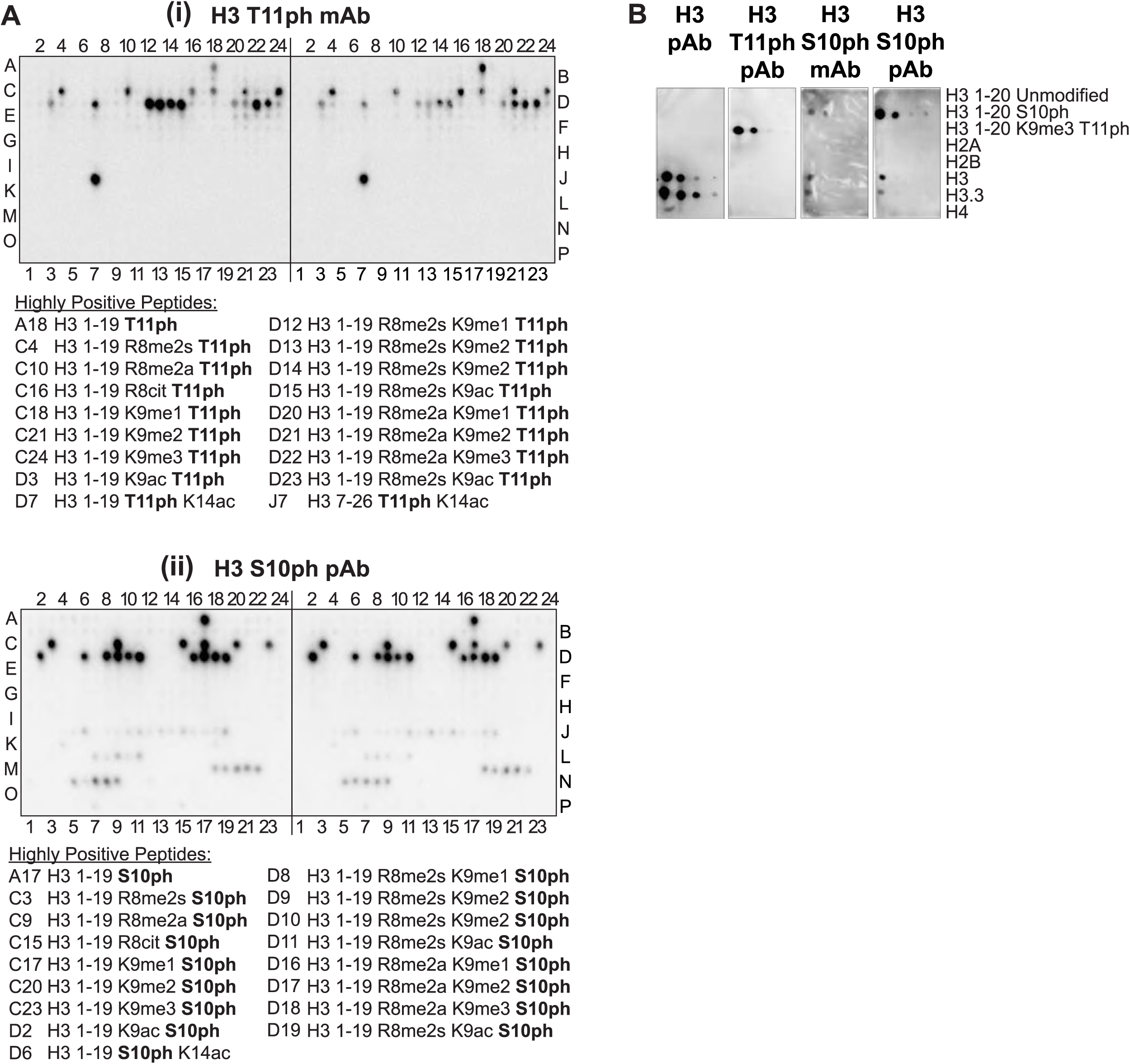
Specificity of anti-H3 T11ph and anti-H3 S10ph antibodies. (**A**) Histone peptide array western blots showing the specificity of (**i**) anti-H3 T11ph mAb or (**ii**) anti-H3 S10ph pAb and their tolerance of neighboring modifications. Blots are of duplicate 384-peptide arrays (MODified Histone Peptide Array, Active Motif 13001) of immobilized synthetic histone H2A, H2B, H3 and H4 unmodified peptides or peptides containing from one to four modified residues including many possible combinations of histone modifications that are found in higher eukaryotes, of which only a small number are known to be present in yeast. Positions A1–L11 contain H3 peptides, L12–O11 contain H4 peptides, O12–P3 contain H2A peptides and P4–P19 contain H2B peptides. Peptides that were highly reactive with either antibody are listed below the blot image; the entire table of peptides is listed in **Supplemental Table S2.** (**B**) Immunodetection of histone H3 amino-terminal peptides (residues 1–20) or recombinant histone proteins spotted onto PVDF membranes demonstrating the specificity of antibodies to phosphoH3 T11 and phospho-H3 S10. Spots were 10-fold serial dilutions of peptides or recombinant histones starting with 167 ng in the left-most column. Recombinant histone proteins produced in *E. coli* were from the following species: H2A, H2B, H3 from *S. cerevisiae*; H3.3 from *H. sapiens*; and H4 from *X. laevis*. Antibodies were: anti-H3 pAb (Abcam 1791), which is specific to the carboxy-terminal 35 amino acids of histone H3; anti-H3 T11ph polyclonal (Active Motif 39151); anti-H3 S10ph monoclonal (EMD Millipore 05-817); and anti-H3 S10ph polyclonal (EMD Millipore 06-560).

***Notes:***

(**Panel Ai**) The diphosphorylated H3 1–19 S10ph T11ph peptide at position D5 was not detected by the anti-H3 T11ph mAb, whereas all phospho-T11 containing peptides (except those that also contained phospho-S10) were detected (peptides containing phospho-T11 along with methyl-K4 were not included in the array). We conclude that this mAb detects only the monophosphorylated peptide, but that it is tolerant of other modifications of the H3 N-terminal tail.

(**Panel Aii**) The diphosphorylated (S10ph T11ph) peptide at position D5 was not detected by the anti-H3 S10ph pAb, whereas all phospho-S10 containing peptides (except those that also contained phospho-T11) were detected (peptides containing phospho-S10 along with methyl-K4 were not included in the array). We conclude that this pAb detects only the monophosphorylated peptide, but that it is relatively tolerant of other modifications of the H3 N-terminal tail. This pAb showed detectable cross-reaction to other modifications as well. Peptides that scored as weakly reactive with anti-H3 S10ph pAb were: J6, H3 1–19 S10ph K14ac; J11, H3 7–26 K18ac; J13, H3 7–26 K14ac R17me2s; J15, H3 7–26 R17me2s K18ac; J19, H3 7–26 K14ac R17me2a K18ac; K4, H3 16–35 S28ph; L7, H3 26–45 unmodified; L8, H3 26–45 K36me1; L9, H3 26–45 K36me2; L11, H3 26–45 K36ac; M18, H4 11–30 unmodified; M19, H4 11–30 K12ac; M20, H4 11–30 K16ac; M21, H4 11–30 R17me2s; M22, H4 11–30 R17me2a; N5, H4 11–30 R24me2a; N6, H4 11–30 R24me2s; N7, H4 11–30 K12ac K16ac; N8, H4 11–30 K16ac R17me2s; N9, H4 11–30 K16ac R17me2a.

(**Panel B**) Anti-H3 T11ph pAb was capable of detecting phospho-T11 even with nearby methylation at lysine 9, a modification that occurs in *S. pombe* and metazoans, but not in *S. cerevisiae*. Both anti-H3 S10ph antibodies also reacted slightly with full-length recombinant H3 and H3.3.

**(A)** *S. pombe* read coverage. The read coverage from a representative sample (H3 ChIP-seq from wild type culture 1 at 3 hr) is shown. The zones of exceptionally high coverage at the ends of chromosome III are the ribosomal DNA repeats. Note that the read density is too sparse to allow investigation of H3 or H3 T11ph patterns across the *S. pombe* genome. **(B)** Reproducibility of read depth for *S. pombe* chromosomes I and II. Stacked bars show the contribution of reads from chromosomes I and II relative to the total for these two chromosomes. Lower case letters are sample identifiers as indicated in **Figure 5D**. Chromosome III had more variable read coverage (data not shown), possibly because of greater sampling error attributable to presence of the large rDNA arrays. Because relative read depths were stable for chromosomes I and II, these were combined to calculate the normalization factors for the *S. cerevisiae* maps. **(C,D)** Reproducibility of spatial patterns for anti-H3 and anti-H3 T11ph ChIP-seq. The map for each wild-type sample (solid lines) is shown separately, superimposed on the *spo11-Y135F* map (dashed lines) for comparison. The maps depict the same region shown in **Figure 5C**. Note that the H3 ChIP-seq maps were highly reproducible across all four wild-type samples as well as the *spo11-Y135F* sample. For H3 T11ph, although the magnitude of the signal varied between wild-type samples, their spatial patterns were highly similar and all gave substantially higher coverage than the *spo11-Y135F* sample.

**(A, B)** ChIP-chip (Panizza *et al.* 2011) and ChIP-seq (this study) yield similar patterns for Red1. In panel A, profiles are shown around the same region on chromosome I as in **Figure 5A–C**. In panel B, the correlation is shown for ChIP-chip and ChIP-seq signals in non-overlapping 5-kb bins genome wide. The good correspondence of the Red1 ChIP-chip and ChIP-seq data validate the use of Hop1 ChIP-chip data (Panizza *et al.* 2011). **(C)** H3 T11ph enrichment around presumed axis-attachment sites. H3 T11ph ChIP-seq coverage is compared with Red1 and Hop1 ChIP-chip data averaged around 1717 Red1 ChIP-seq peaks. The green lines in the lower graph show the ratio of H3 T11ph to H3 ChIP-seq. Compare with **Figure 5E**. **(D)** H3 T11ph correlates well with Red1 and Hop1 genome wide. Each point compares the H3 T11ph ChIP-seq coverage in wild type with Red1 or Hop1 ChIP-chip signal (Panizza *et al.* 2011) averaged across non-overlapping 5-kb bins. Compare with **Figure 5F**. Correlation coefficients (Pearson’s *r*) are indicated in each plot. **(E)** H3 T11ph and Red1 around DSB hotspots. Patterns around Spo11-oligo hotspots are plotted as in **Figure 5G**, except that Red1 ChIP-chip data are used.

**Figure S2.**
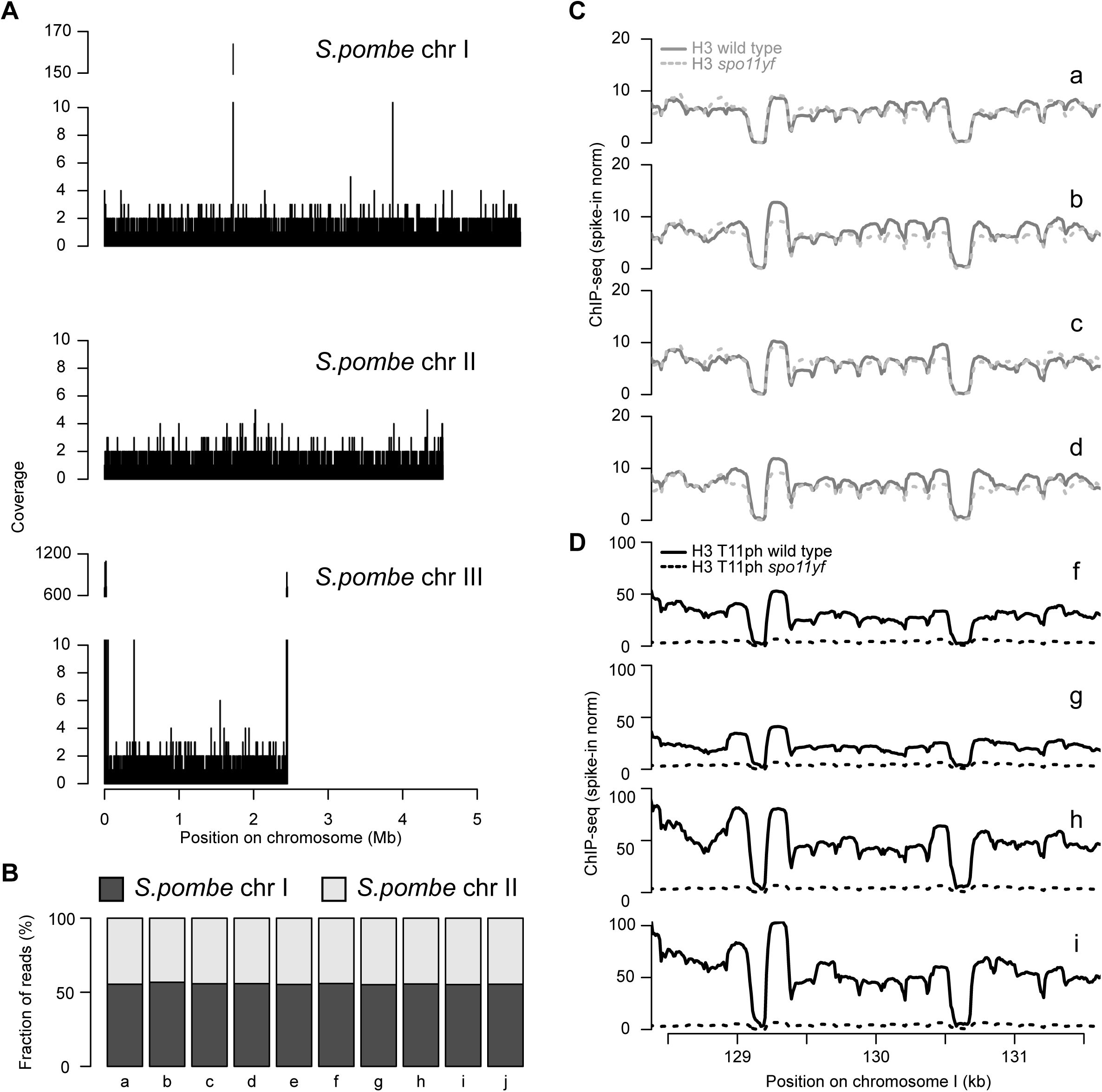
Anti-H3 and anti-H3 T11ph ChIP-seq.

**Figure S3.**
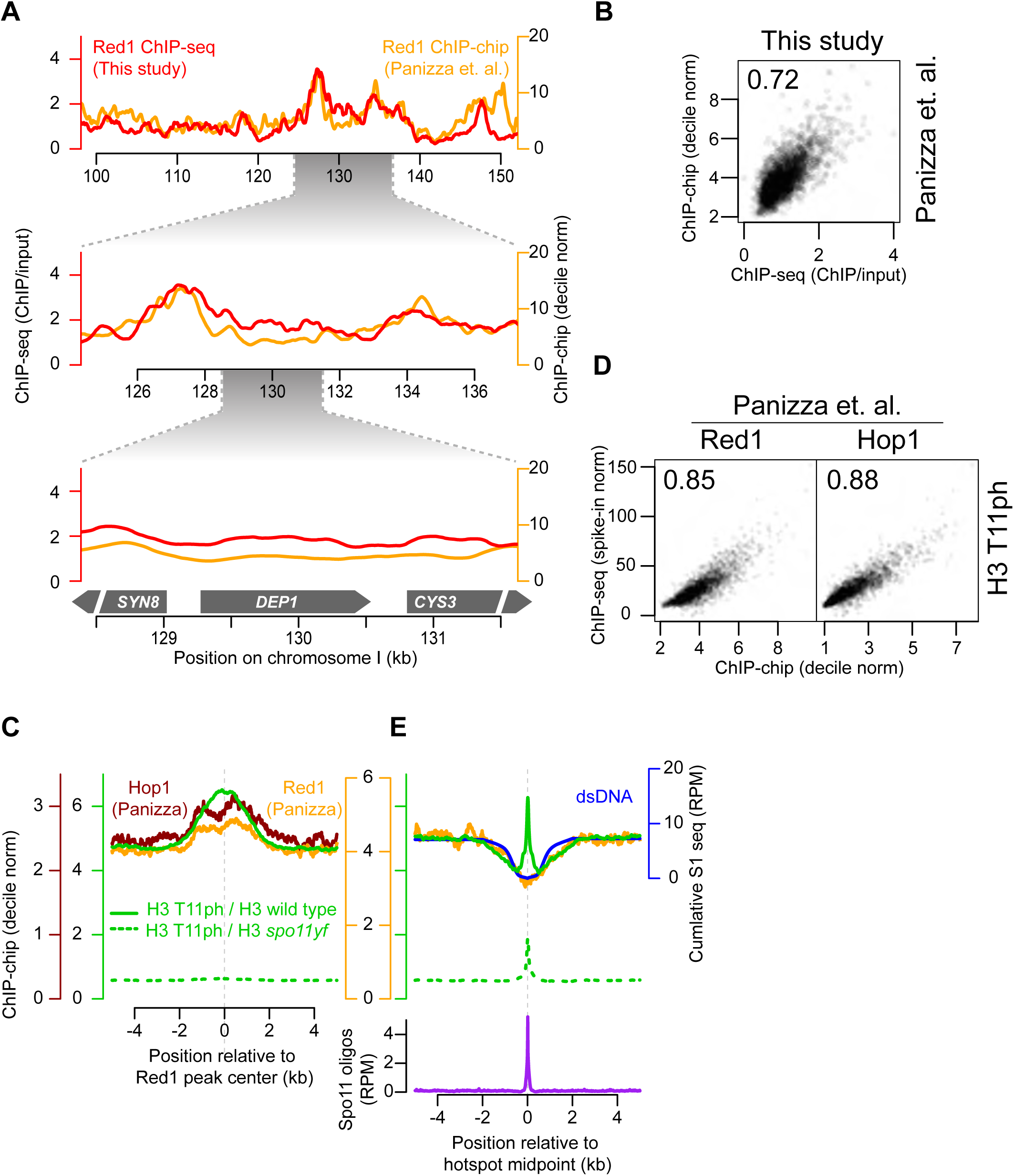
Further comparisons of H3 T11ph with Red1 and Hop1 ChIP

**Supplemental Table S1.**
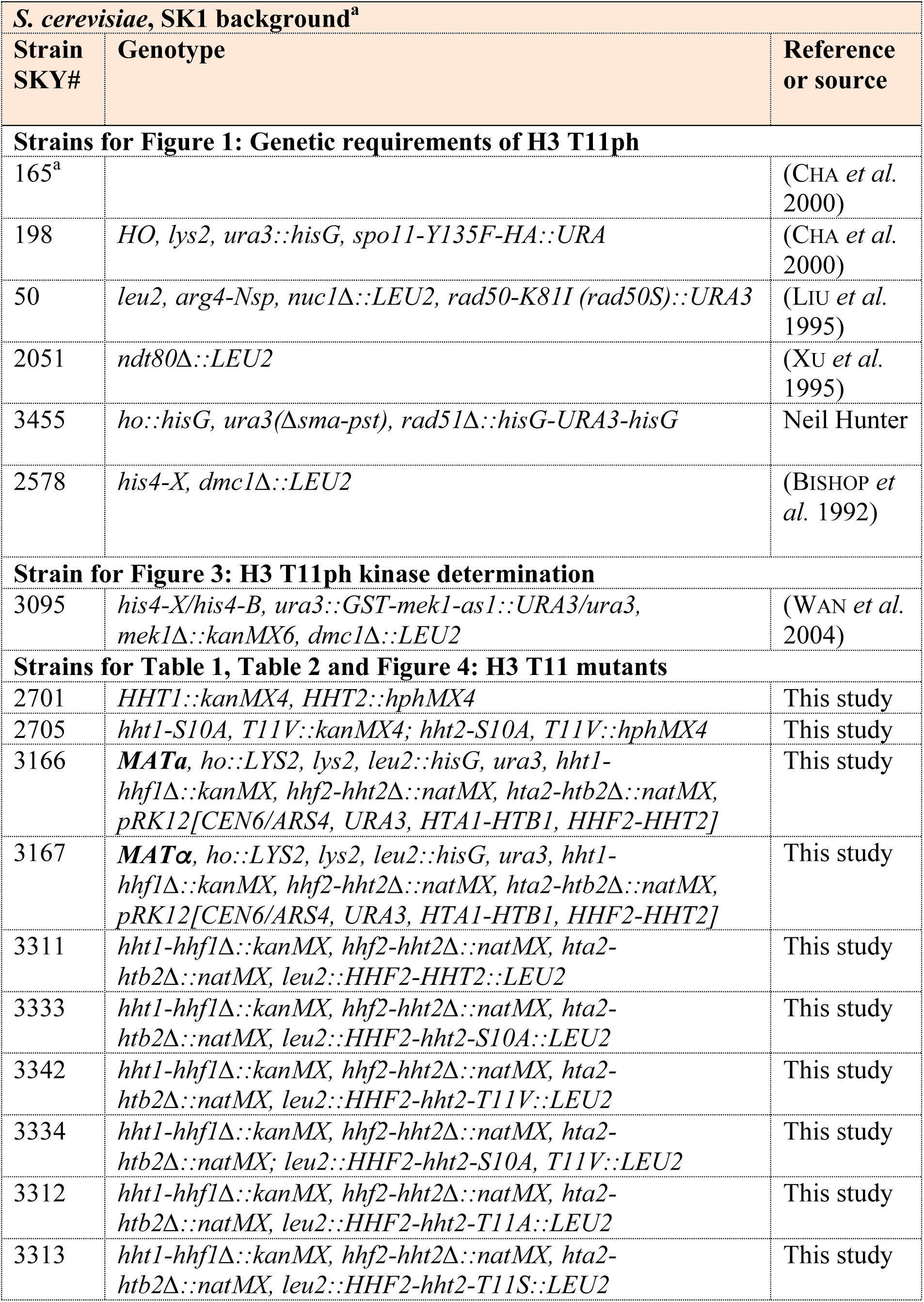
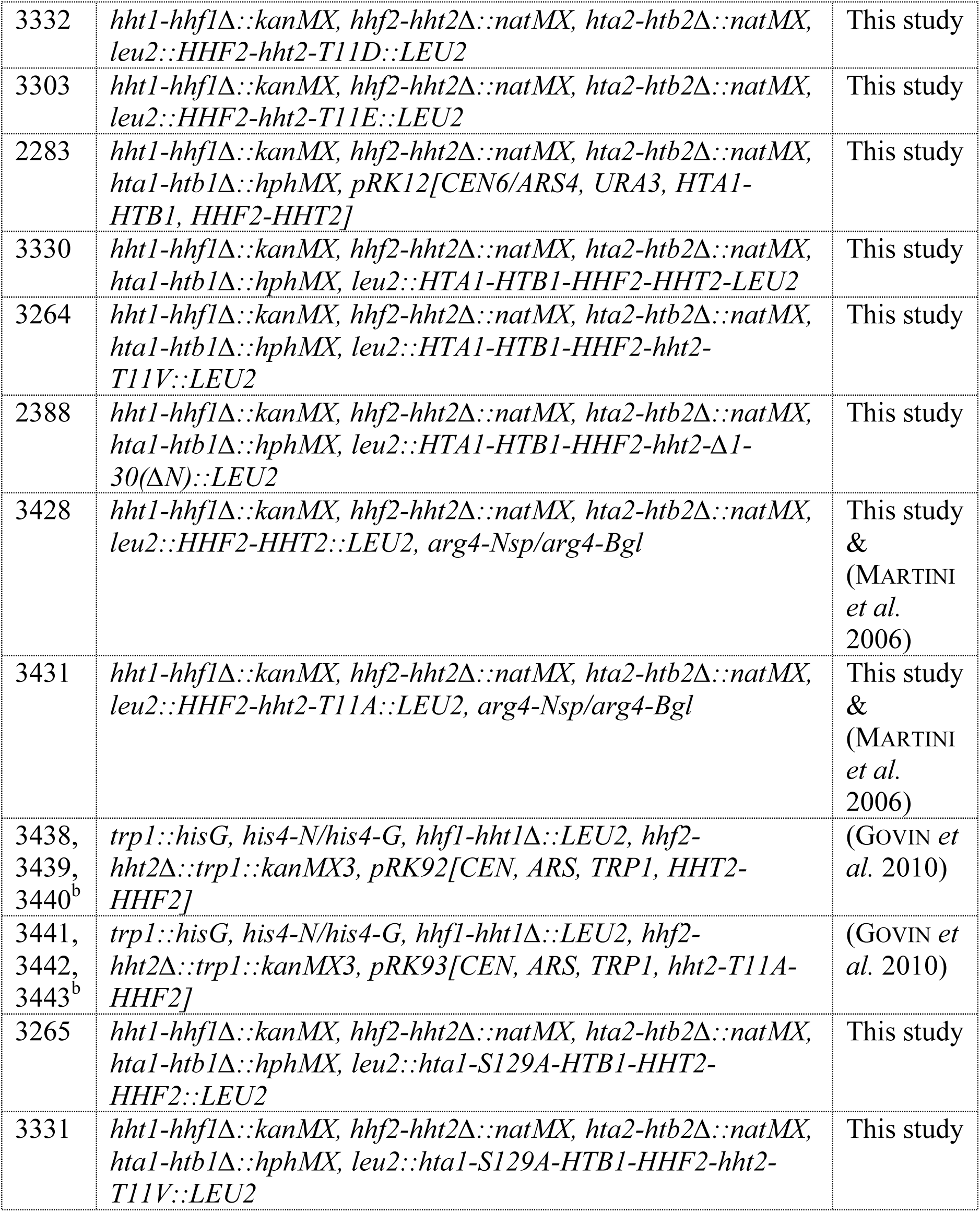
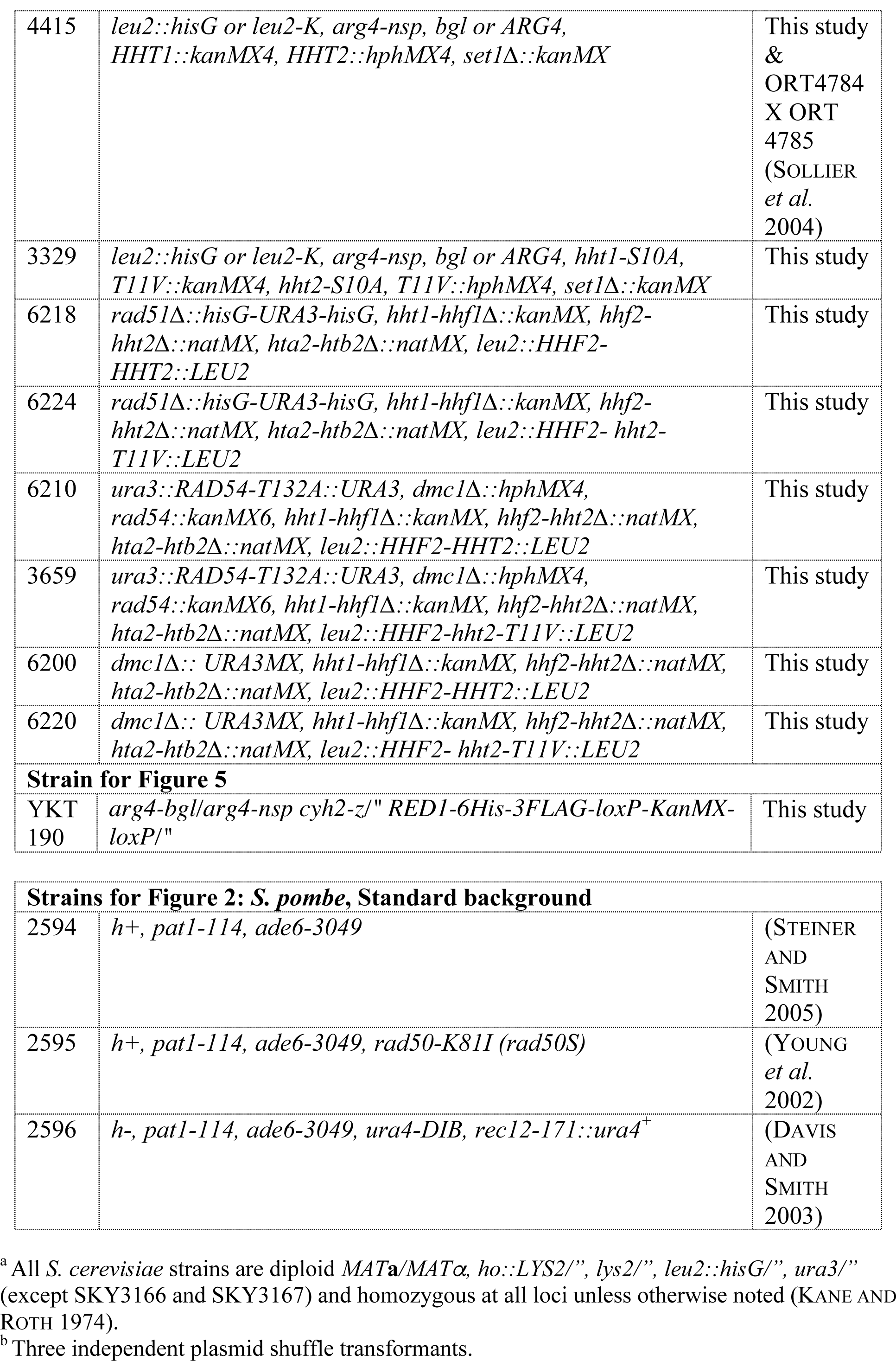
List of *S. cerevisiae* and *S. pombe* strains used in this study.

**Supplemental Table S2.**
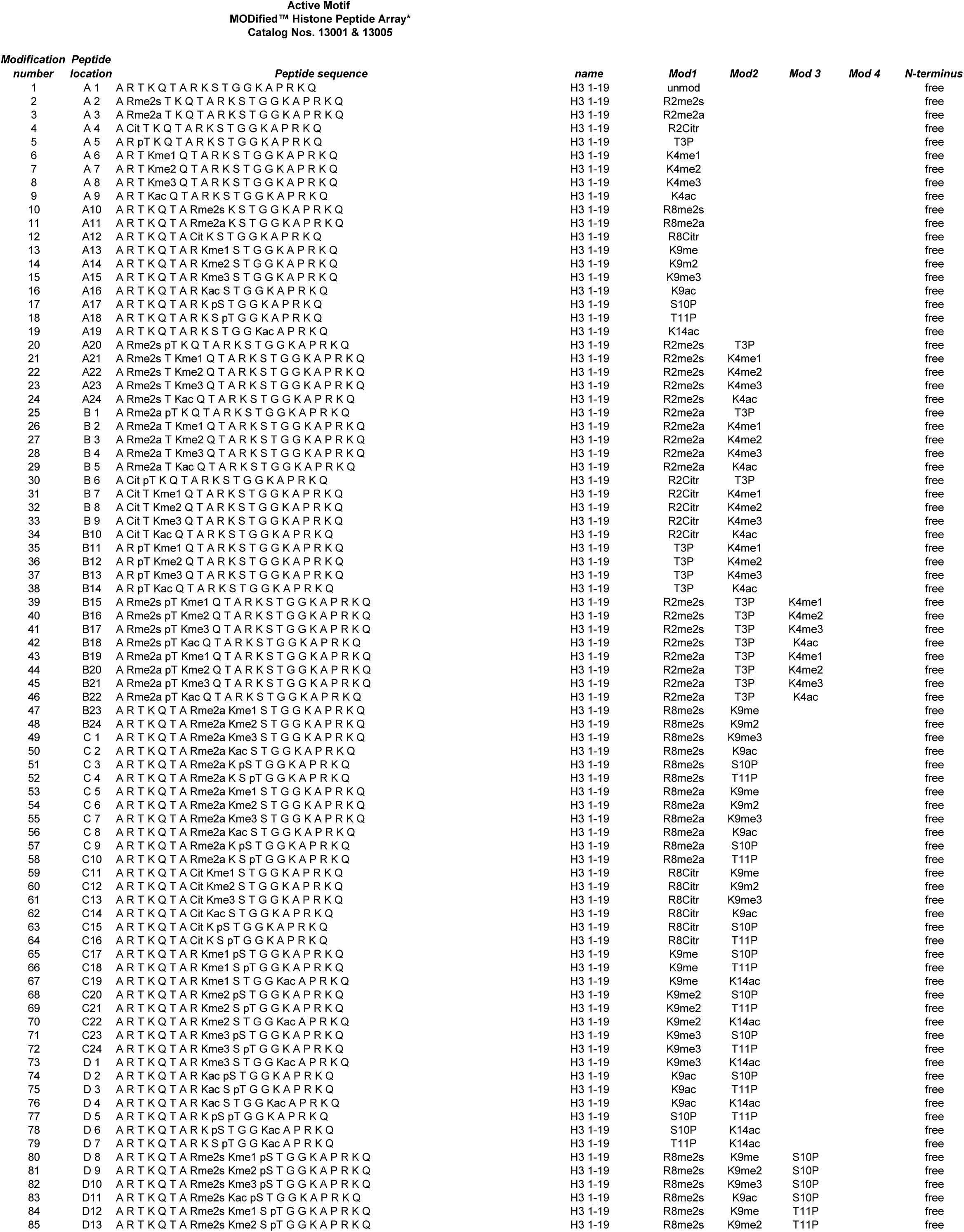
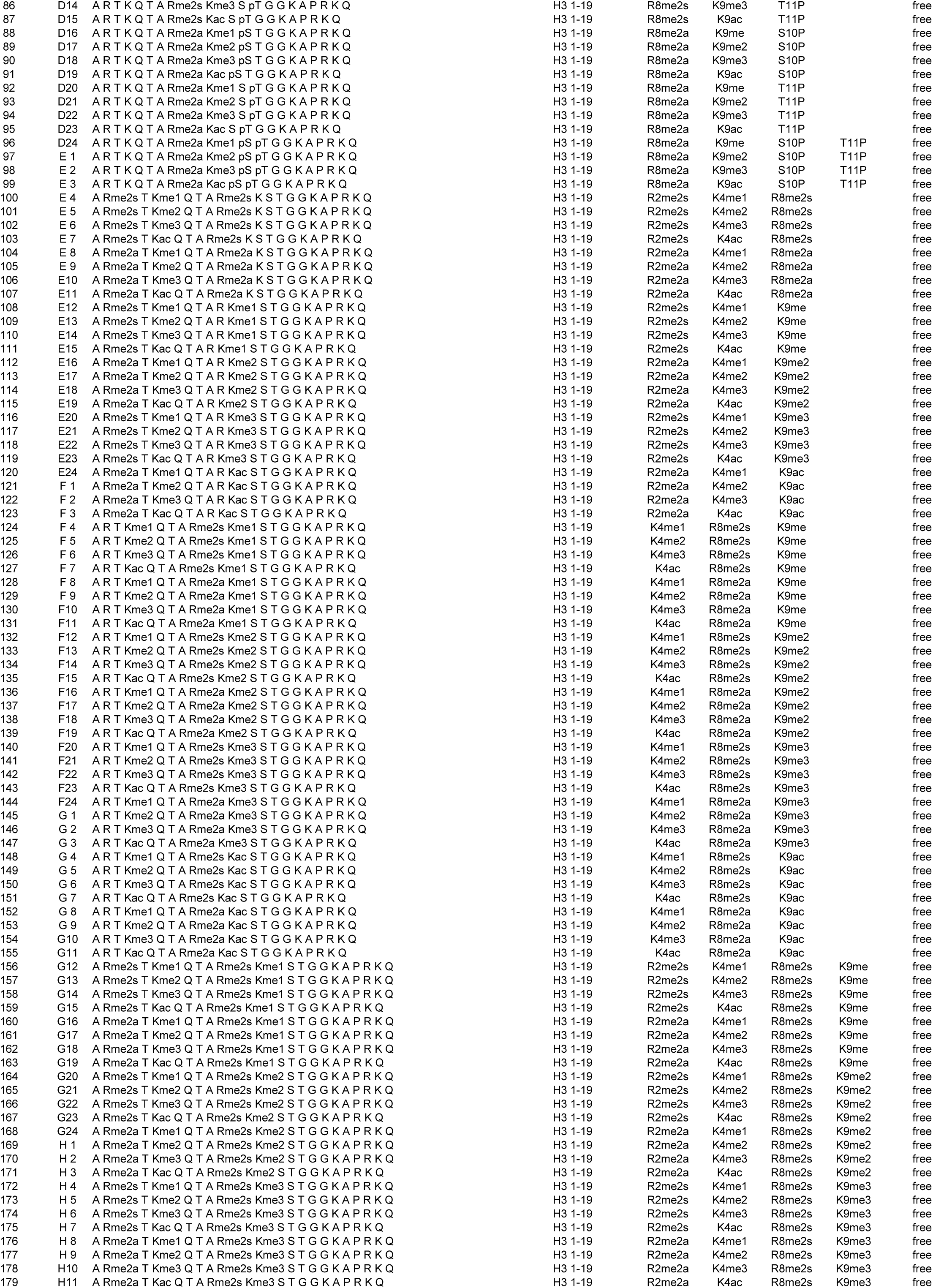
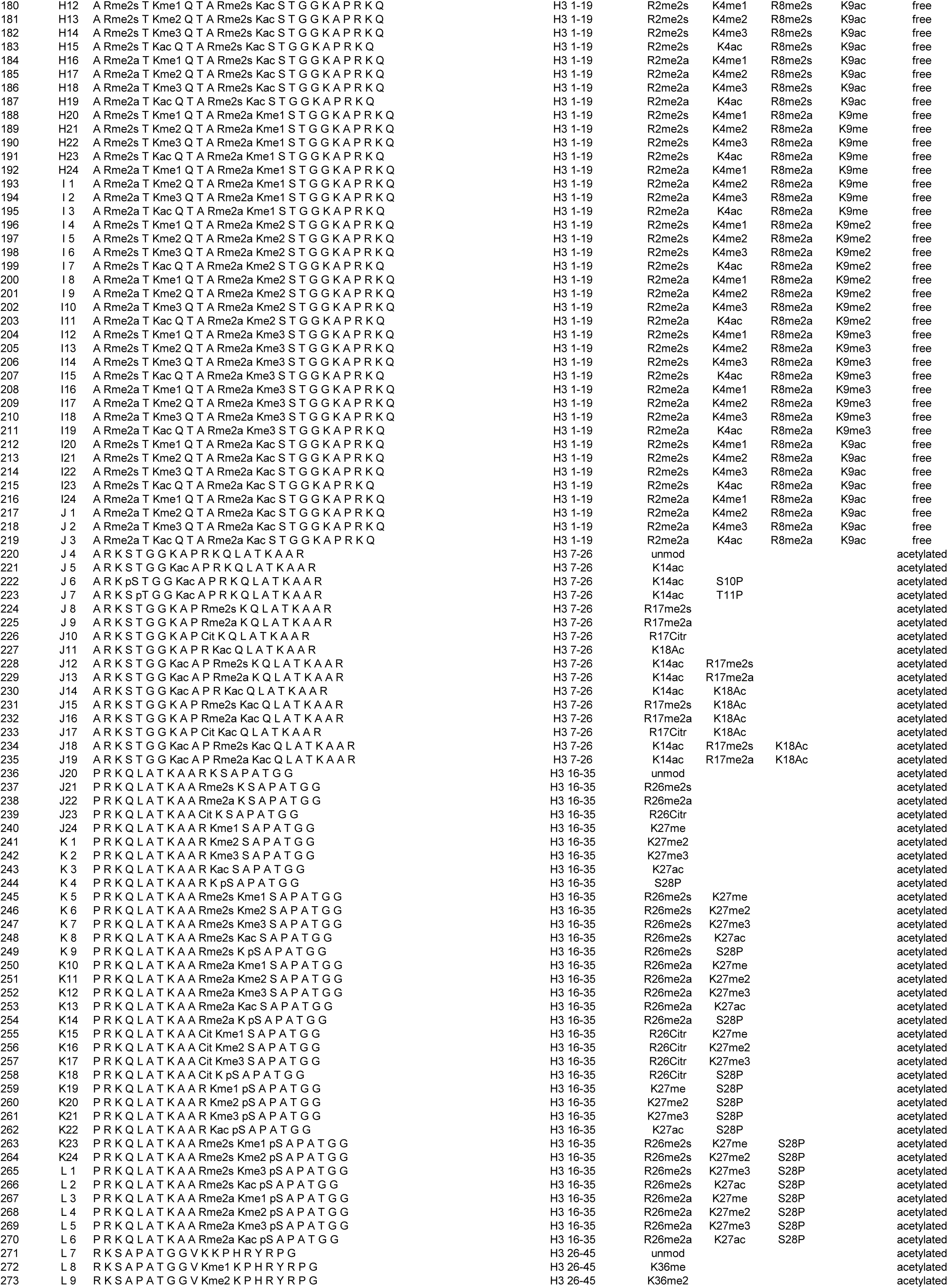
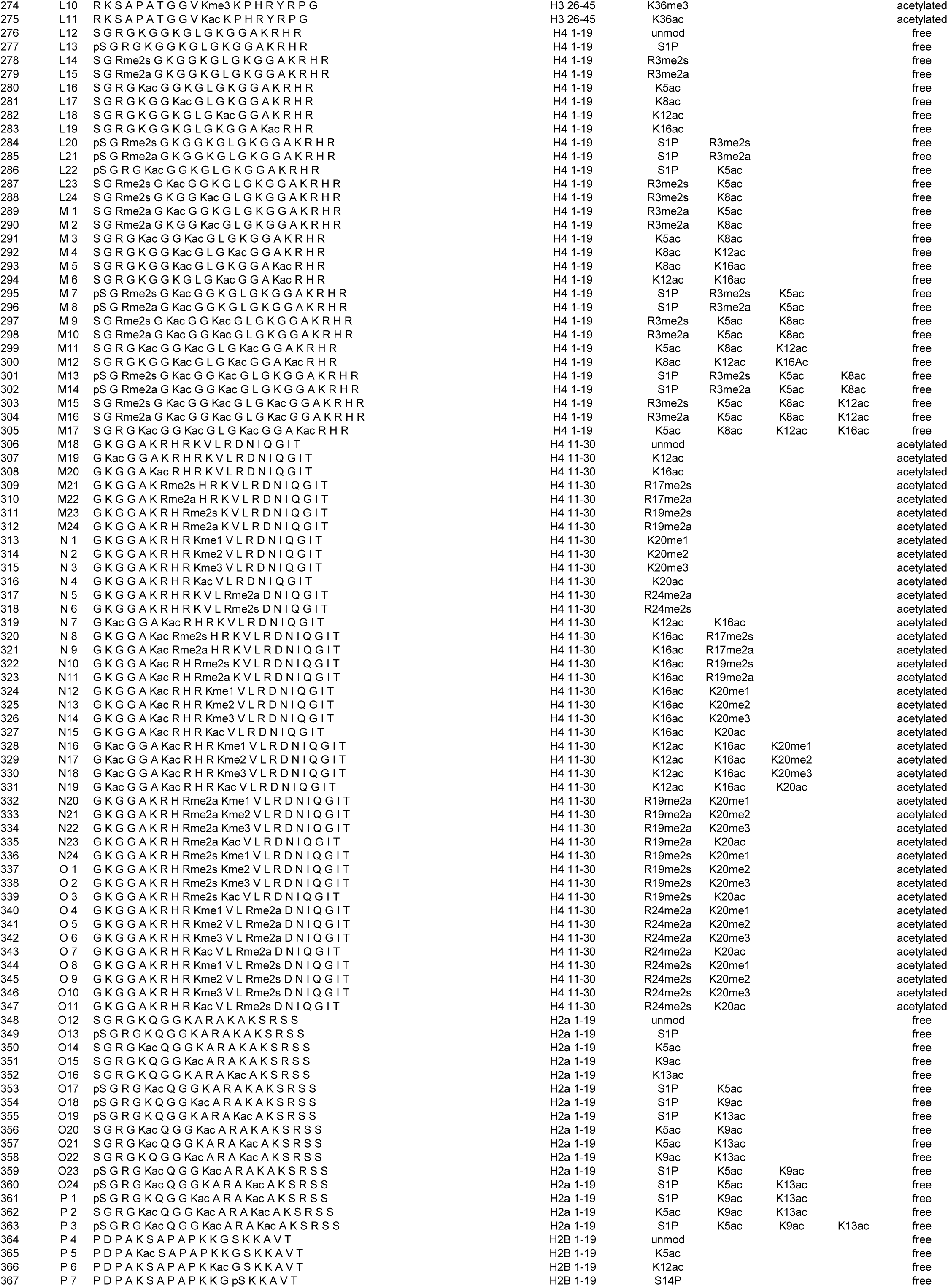
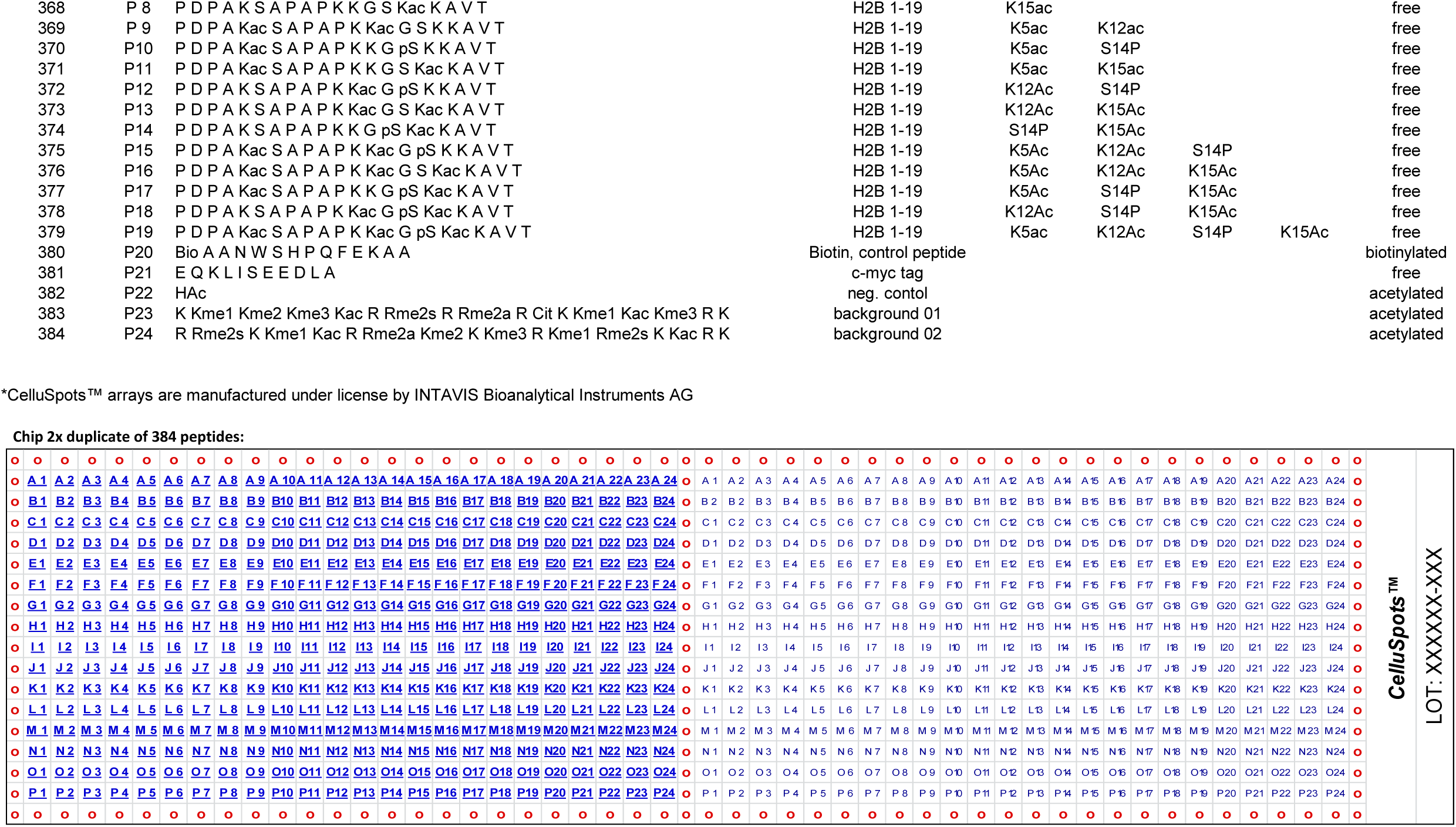
Synthetic peptide array.

